# Elimination of reference mapping bias reveals robust immune related allele-specific expression in crossbred sheep

**DOI:** 10.1101/619122

**Authors:** Mazdak Salavati, Stephen J. Bush, Sergio Palma-Vera, Mary E. B. McCulloch, David A. Hume, Emily L. Clark

## Abstract

Pervasive allelic variation at both gene and single nucleotide level (SNV) between individuals is commonly associated with complex traits in humans and animals. Allele-specific expression (ASE) analysis, using RNA-Seq, can provide a detailed annotation of allelic imbalance and infer the existence of cis-acting transcriptional regulation. However, variant detection in RNA-Seq data is compromised by biased mapping of reads to the reference DNA sequence. In this manuscript we describe an unbiased standardised computational pipeline for allele-specific expression analysis using RNA-Seq data, which we have adapted and developed using tools available under open licence. The analysis pipeline we present is designed to minimise reference bias while providing accurate profiling of allele-specific expression across tissues and cell types. Using this methodology, we were able to profile pervasive allelic imbalance across tissues and cell types, at both the gene and SNV level, in Texel x Scottish Blackface sheep, using the sheep gene expression atlas dataset. ASE profiles were pervasive in each sheep and across all tissue types investigated. However, ASE profiles shared across tissues were limited and instead they tended to be highly tissue-specific. These tissue-specific ASE profiles may underlie the expression of economically important traits and could be utilized as weighted SNVs, for example, to improve the accuracy of genomic selection in breeding programmes for sheep. An additional benefit of the pipeline is that it does not require parental genotypes and can therefore be applied to other RNA-Seq datasets for livestock, including those available on the Functional Annotation of Animal Genomes (FAANG) data portal. This study is the first global characterisation of moderate to extreme ASE in tissues and cell types from sheep. We have applied a robust methodology for ASE profiling, to provide both a novel analysis of the multi-dimensional sheep gene expression atlas dataset, and a foundation for identifying the regulatory and expressed elements of the genome that are driving complex traits in livestock.

## Introduction

Allele-specific expression (ASE) is the imbalance of allelic expression between parental (diploid) copies at the same locus (Barlow and Bartolomei, 2014). It is most commonly associated with cis-acting regulatory variation that may mediate parent-of-origin, sex- or tissue-specific transcription of one allele relative to the other (Renfree et al., 2009; Hasin-Brumshtein et al., 2014). In a single individual, where there are informative sequence variants (i.e. heterozygote loci) that distinguish the products of two alleles, ASE can be detected by RNA-sequencing (Chamberlain et al., 2015; GTEx Consortium et al., 2017; Cao et al., 2019; Guillocheau et al., 2019). The ratio of allelic read counts obtained from RNA-Seq datasets can be used as a reliable proxy for ASE (i.e. *ASEratio* = *CountsAllele*1/(*CountsAllele*1 + *CountsAllele*2)) (Edsgärd et al., 2016).

Large and complex RNA-Seq datasets give rise to unique and interesting computational challenges, in particular the elimination of reference mapping bias in ASE analysis of diploid genomes. RNA-Seq data are commonly mapped against reference genomes which are typically ‘flat’, with each position represented only by the reference (most abundant) allele. As such, reads containing heterozygous loci are more likely to be erroneously mapped (Degner et al., 2009; Stevenson et al., 2013; Hodgkinson et al., 2016). This can lead to high false positive ASE locus discovery rates (Degner et al., 2009). Although development of de novo transcript assemblers (Zerbino and Birney, 2008), usage of personalised reference genomes (Rozowsky et al., 2011; Smith et al., 2013), variant-aware aligners (Xin et al., 2013; Hach et al., 2014) and mapping-free quantification e.g. Kallisto (Bray et al., 2016) have resolved some of these issues, reference allele mapping bias remains a considerable challenge in ASE studies. In the absence of ‘trios’ of animals or reference population phased haplotype information, which are rare for livestock, correction of mapping bias via synthetic reads with either N masking or alternative mapping bias correction at the heterozygote sites, has proven a robust alternative for ASE discovery (Degner et al., 2009; Mayba et al., 2014; van de Geijn et al., 2015; Miao et al., 2018). In 2015 Van de Geijn et al. benchmarked the WASP software mapping correction strategy against N-masked reads and personal genome mapping. WASP showed consistent correct mapping of reads with multiple alleles and lower false discovery rates in comparison to the other two methods (van de Geijn et al., 2015). The analysis pipeline we present in this manuscript is based on WASP’s methodology and is designed to minimise reference bias while providing accurate profiling of allele-specific expression in large and complex RNA-Seq datasets.

We have developed an ASE analysis pipeline using the combination of software available under open licence, WASP (reference mapping bias removal) (van de Geijn et al., 2015), GATK (ASEReadCounter) (McKenna et al., 2010; Van der Auwera et al., 2013) and GeneiASE (Liptak-Stouffer aggregative ASE gene model) (Edsgärd et al., 2016). The GeneiASE model is capable of testing ASE at the gene level using two approaches: i) Static ASE which measures allelic imbalance within a gene (i.e. when ASE variants are located within the boundaries of the gene) and ii) Individual condition dependent ASE (ICD) which measures inducible ASE in a gene under an environmental pressure between 2 timepoints (i.e. in stimulated or unstimulated immune cells).

In addition to ASE at the gene level, we can also measure significant ASE at the single nucleotide level (SNV). ASE has been shown to be enriched within expression quantitative trait loci (eQTL) regions (Montgomery et al., 2010), therefore identifying ASE variants can be useful for understanding the transcriptomic control of complex traits in livestock. Complex trait mapping of ASE loci has been associated with phenotypes such as resistance to Marek’s disease in chicken (Meydan et al., 2011) and pigmentation patterns in sheep (García-Gámez et al., 2011).

Understanding ASE is also important because cross-breeding now underlies most livestock production systems. Knowledge of ASE may provide insights into the molecular basis of the complex phenomenon of hybrid vigour, as emphasised by recent studies on two Chinese goat breeds and their F1 hybrids (Cao et al., 2019) and in F1 crosses of two highly inbred chicken lines (Zhuo et al., 2017). In this study we measure ASE in crossbred sheep. Sheep are an economically important livestock species in many countries across the globe and particularly in emerging economies. The identification of prevalent ASE in populations or breeds, especially in economically relevant phenotypes and tissues, could be used to improve genomic prediction in sheep breeding programmes, such as those that have been established in Australia and New Zealand (Daetwyler et al., 2010).

Using the methodology we describe, for mapping bias correction and robust positive ASE discovery, we were able to profile pervasive allelic imbalance across tissues and cell types, at both the gene and SNV level, in Texel x Scottish Blackface sheep. We analysed a subset of total RNA-Seq libraries from liver, spleen, ileum, thymus and bone marrow-derived macrophages (BMDM) (+/−) lipopolysaccharide (LPS), from six individual adult crossbred sheep to produce a detailed picture of allelic imbalance in immune-related tissues and cell types. We chose to focus this analysis on immune related tissues in part because of the depth of available sequence in those tissues, and in part because they contain abundant immune cell populations. The diversity of cell populations is reflected in the transcriptional complexity of immune tissues and cell types in the sheep gene expression atlas dataset (Clark et al., 2017; Bush et al., 2018). As such this subset of tissues gave us a transcriptionally rich dataset in which to measure ASE. We also included BMDMs stimulated and unstimulated with LPS to mimic infection with Gram-negative bacteria, in order to test whether ASE changed in response to stimulation with LPS in these cells. By measuring ASE in these tissues and cell types from sheep we were able to: i) provide insight into how pervasive ASE is across tissues at the gene and SNV level, ii) generate tissue-specific ASE profiles, iii) investigate sex-specific patterns of ASE, and iv) determine the extent to which ASE changes in response to stimulation with LPS in an immune cell type. This novel analysis of the multi-dimensional sheep gene expression atlas dataset provides a foundation for further analysis of the regulatory and expressed elements of the genome that are driving complex traits in sheep.

## Methods

### Sample preparation and RNA extraction

Data from three male and three female Texel x Scottish Blackface (TxBF) sheep from the sheep gene expression atlas project (Clark et al., 2017) were used in this study. The dataset including: 1 cell type (BMDMs (+/−) LPS treatment) and 4 tissues (thymus, spleen, liver and ileum). Tissue collection, storage and RNA extraction is described in (Clark et al., 2017). BMDM’s were cultured in vitro for 7 days in the presence of macrophage colony-stimulating factor (CSF1 (10^4^ U/ml)) and unstimulated (0 hrs - LPS) and stimulated (7hrs +100ng/ml LPS) samples of BMDMs were obtained also as described in (Clark et al., 2017). A total of 2 samples (1 thymus and 1 spleen) did not pass the RNA quality control (RNA integrity number (Mueller et al., 2004); RIN >7) and were not included in the sheep gene expression atlas. Library preparation was performed by Edinburgh Genomics (Edinburgh Genomics, Edinburgh, UK). All total RNA Illumina TruSeq libraries (125bp paired end) were sequenced at a depth of > 100 million reads per sample.

### Reference mapping bias removal

BAM files from RNA-Seq data, were previously produced by mapping fastq files to the Oar v3.1 top level DNA fasta track, using HISAT2 (default mismatch penalty MX=6 MN=2) as described in (Clark et al., 2017). Detailed settings and parameters for all the tools used to generate the BAM files can be found at (FAANG, 2018). These BAM files were used to locate reads with heterozygote loci using WASP’s find_intersecting.py script (van de Geijn et al., 2015). The intersection of reads and heterozygote loci in all samples were based on the Ensembl v92 variant call format (VCF) track (Ensembl v92: ovis_aries_incl_consequences.vcf.gz). Briefly the Ensembl VCF file was filtered for bi-allelic variants within exonic regions, 5k up or downstream of exonic regions (5’ or 3’ UTRs) and intronic regions of all transcripts within the Oar3.1 sheep assembly (exclusion of indels and intergenic variants). These variants were used in WASP’s find_intersecting.py script in order to extract reads mapped to coordinates containing variants for each gene. As a result, reads aligned to exonic, 5’ or 3’ UTRs and intronic regions were separated into reads intersecting heterozygote loci and reads that did not intersect heterozygote loci. Synthetic copies of reads intersecting heterozygote loci, were created with the alternate allele flipped to the remaining options of A, T, C or G (up to 6 loci/read[2^n^] max 64 combinations of synthetic reads) using parameters defined in WASP (van de Geijn et al., 2015). This was followed by remapping of the synthetic reads using HISAT2 (default mismatch penalty MX=6 MN=2) (Li and Durbin, 2009; Kim et al., 2015) and eliminating the original reads (and their synthetic copies) which mapped to a different coordinate in any of its synthetic copies (WASP’s filter_reads.py) (van de Geijn et al., 2015). After merging the retained reads with that did not intersect heterozygote loci, a final BAM file was produced for ASE read counting step (WASP’s remove_dup.py).

### Allelic read counts and depth filtration

Allele-specific read counting was carried out using the ASEReadCounter module of GATK v3.8 with parameters -mmq 50 -mbq 25 (McKenna et al., 2010). Multiple pre-processing steps were performed prior to GeneiASE input as instructed by (Edsgärd et al., 2016) which included, preparing per chromosome indices, merging the variant set with corresponding gene coordinates and bi-allelic expression filtering. Loci with < 10 reads mapped were excluded, as were loci with < 3 reads, or < 1% of the total reads, mapped to both the reference and alternative allele. This form of filtration will eliminate loci exhibiting mono-allelic expression (MAE) as previously described (Degner et al., 2009; Stevenson et al., 2013; Mayba et al., 2014). Producing evidence of MAE using total RNA-Seq datasets produced by Illumina short read sequences without parent of origin genotypes or imprinting information has been a controversial issue (DeVeale et al., 2012). Our dataset did not include the trios of animals or personalised genomes that would be necessary to resolve MAE. As such we decided to exclude MAE altogether for our analysis using stringent bi-allelic filtration criteria. Similar bi-allelic filtration criteria have been previously used routinely in ASE studies (Mayba et al., 2014; Chen et al., 2016a; Edsgärd et al., 2016; GTEx Consortium et al., 2017; Raghupathy et al., 2018; Cao et al., 2019; Guillocheau et al., 2019; Gutierrez-Arcelus et al., 2019). The workflow of the analysis pipeline for ASE analysis is detailed in Fig.1.

**Figure 1.**
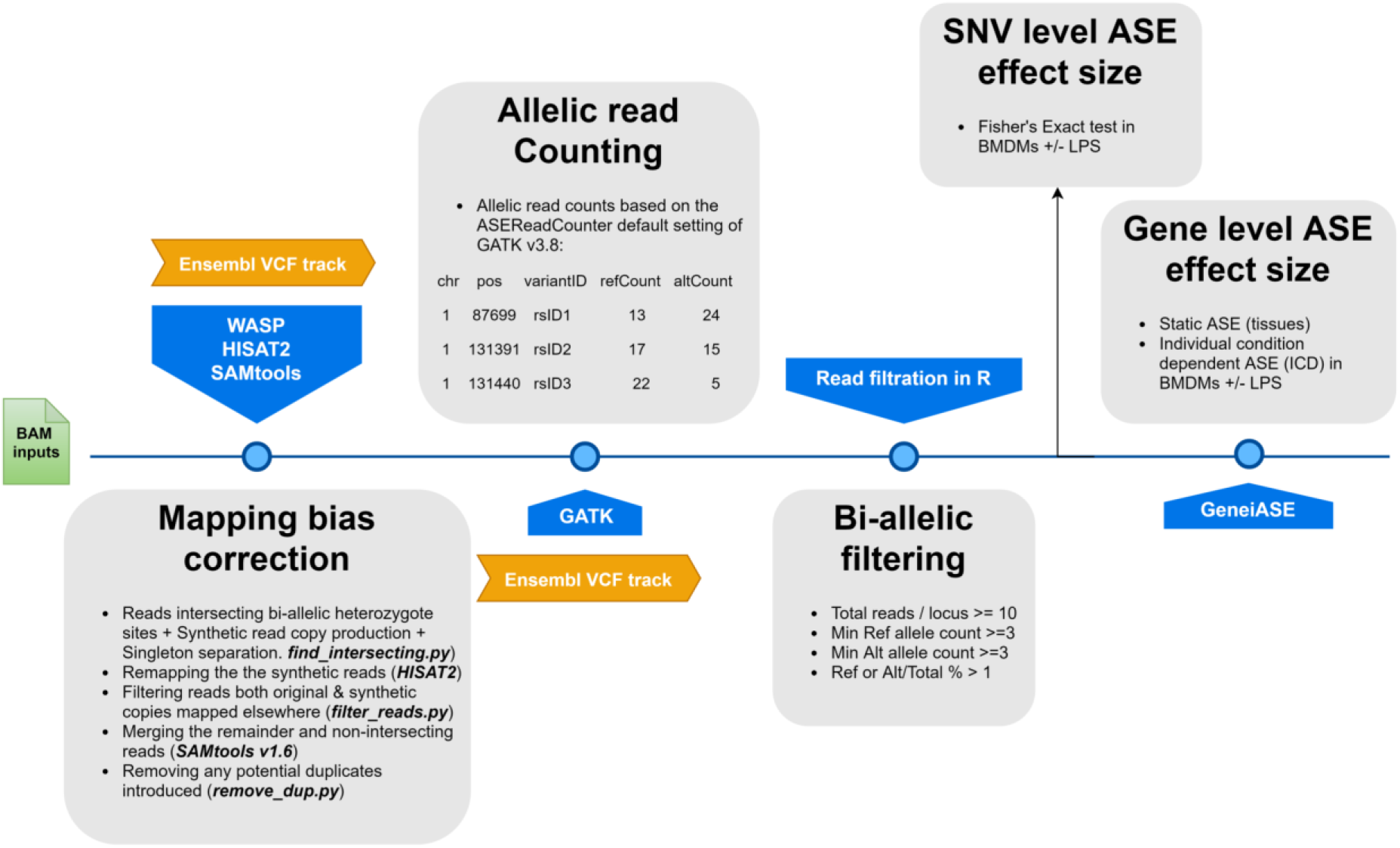
A flowchart of the allele-specific expression analysis pipeline applied to the sheep gene expression atlas dataset and optimised for WASP and GeneiASE programmes. The remapping was carried out using HISAT2 (Kim et al., 2015) in combination with SAMtools (Li et al., 2009). The Genome Analysis Toolkit v 3.8 was used for the ASE read counting section.

### Experimental design for defining allele-specific expression

ASE was defined according to the following three categories:

i. Static ASE: which is inherent allelic imbalance (AI) in each gene calculated by ASE at all heterozygote loci (i.e. *ASE* = *Counts RefAllele*/(*Counts RefAllele* + *Counts AltAllele*) within the boundaries of the gene. The effect size of ASE at gene level was produced by aggregation of the ASE effect size at SNVs within the gene boundaries (the Liptak-Stouffer method, applied by the GeneiASE aggregative model). A null distribution of ASE effect size for genes in each transcriptome was produced by random sub-sampling (n = 1e+5) from a pool of genes having min 2 and max 100 loci within their boundaries. The ASE effect size of each gene (aggregated using Liptak-Stouffer) was then tested against the null distribution of same SNV number, via a modified bi-nomial test (2×1 table). Distribution of p values was examined for uniformity prior to false discovery rate correction (Supplementary Fig.S1, S2 and S3) (Benjamini and Hochberg, 1995; Pounds and Cheng, 2006; Barton et al., 2013).
ii. Individual condition-dependent ASE (ICD-ASE): in which the same ASE effect size was calculated for each gene in the treated versus the untreated timepoints of the same sample (i.e. BMDM +/− LPS). The log2ratio (ASE_treated_/ASE_untreated_) was used in a beta-binomial test (2×2 table) similar to the static mode. The details of this aggregative model have been previously described in the GeneiASE publication (Edsgärd et al., 2016).
iii. Condition dependent ASE at SNV level: in which a contingency table was produced for read counts (ref and alt) for every SNV, present both in treated and untreated conditions (BMDM +/− LPS) (2×2 table) and a Fisher’s exact test performed followed by p value multiple testing correction (Benjamini and Hochberg, 1995). The p values from loci showing ASE and shared by the six adult sheep (ID and coordinate) were unified, using the Stouffer method (Dewey, 2016, 2018) and presented as false discovery rate (FDR) for each locus.

Static ASE was calculated in both tissues and BMDMs (each timepoint was considered separately for BMDMs). Condition dependent ASE analysis was carried out only in BMDMs ± LPS both at gene (ICD-ASE) and SNV (Fisher’s exact) level to study LPS inducible ASE.

### Statistical analysis and thresholds applied

The extraction, transformation and loading of the all datasets and subsequent statistical analysis was carried out in R v ≥ 3.4 unless stated otherwise (R Core Team, 2017). System query language join statements (Wickham et al., 2019) were used to compare lists of ASE genes or SNVs between samples. Raw p values resulting from all three types of ASE analysis were corrected for multiple testing via Benjamini-Hochberg false discovery rate calculations (Benjamini and Hochberg, 1995). The passing threshold of significance in all analyses was considered to be FDR <0.1 (10%) except for the Fisher’s exact test association study. Genes showing ASE in multiple tissues were considered those for which 4 or more of the 6 sheep had significant ASE.

## Results

### Estimation of heterozygous sites across all individuals

To determine the level of heterozygosity present in the RNA-Seq data we first assessed the number of bi-allelic heterozygote sites per individual for each of the six sheep (range= 5,673,703-6,438,497) detailed in Fig.2. Individual variation was observed in the SNVs per gene in each sheep (Fig.2A&C). However, there was no significant difference in the total number of bi-allelic SNVs captured in the RNA-Seq data across all six individuals or between the male and female sheep included in the study (Fig.2B). The bi-allelic SNVs captured in the RNA-Seq data set were annotated using the Ensembl v.92 (Zerbino et al., 2018) reference VCF track. The distribution of SNVs per gene in the Ensembl track is tail-inflated in comparison to the RNA-Seq data Fig.2A. This issue could be due to erroneous assignment of SNVs in hypervariable and repetitive regions, multi-allelic SNVs or simply that there are variants in the Ensembl track that are not expressed (transcribed). The distribution of SNVs for each individual is shown in Fig.2C.

**Figure 2.**
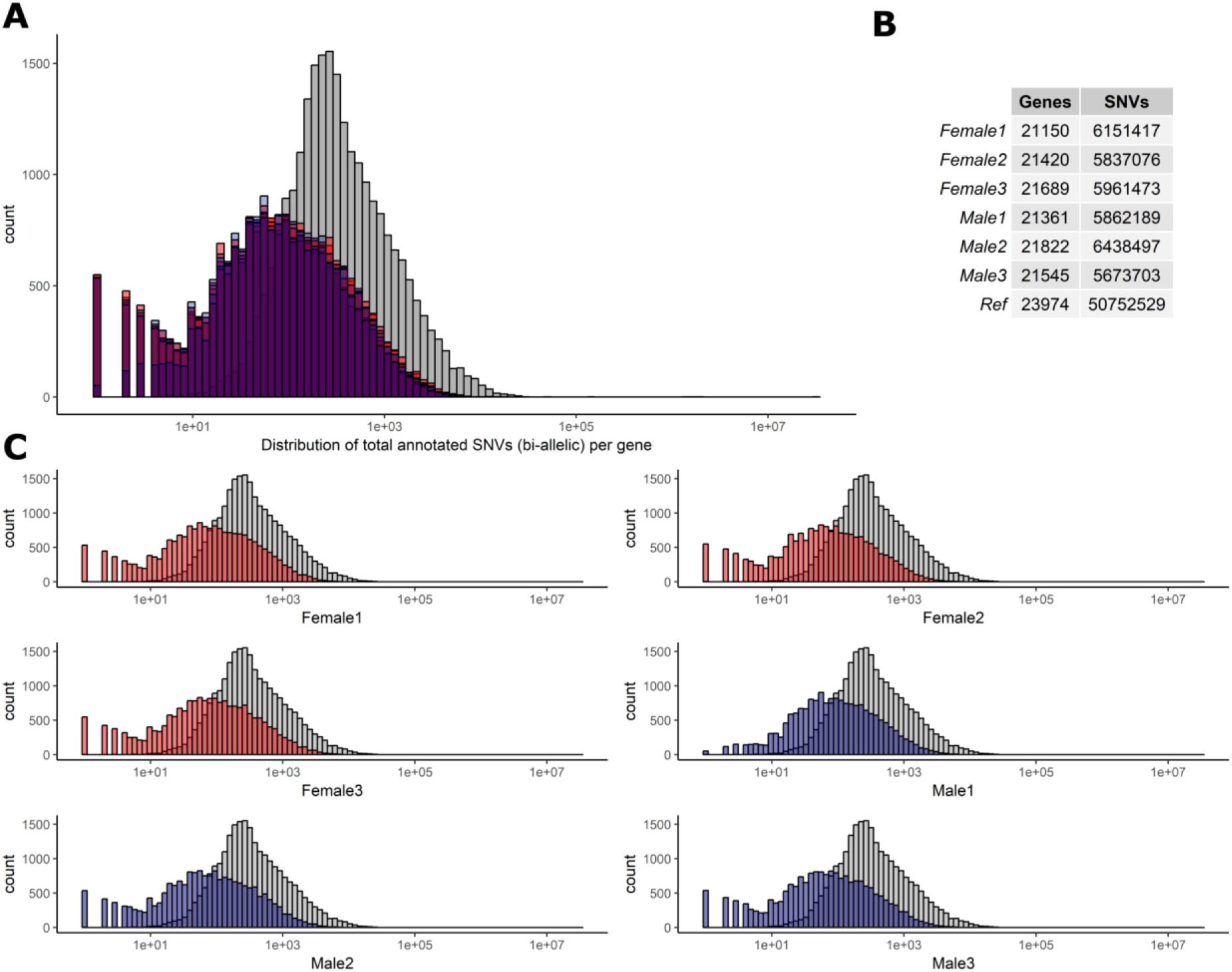
Distribution of biallelic SNVs expressed per gene in each of the 6 TxBF sheep. The total number of SNVs was averaged across thymus, liver, ileum and spleen for every animal. Over 5e+07 SNVs were gathered using Ensembl v.92 VCF track. The total number of SNVs per genes is averaged across 4 tissue RNA-Seq in each animal (~5.9e+06). **A)** Histogram of SNVs per gene counts in the reference track (Ensembl in grey) and 6 sheep in red (females) and blue (males) overlaid. **B)**. The overall numbers of genes and SNVs detected in each animal (averaged over 4 tissues). **C)** Individual histograms from section A with females in red and males in blue.

### Reference mapping bias elimination and quality control

We used the WASP ref bias removal script to successfully minimise ref allele mapping bias in the RNA-Seq samples. The mapping bias was assessed by global distribution of the allelic ratio i.e. ref_counts_/alt_counts_ + ref_counts_ in each RNA-Seq sample as shown in Fig.3 (WASP metrics are included in Supplementary Fig.S10, S11 and S12). The ASE discovery rate at the SNV level on average constituted 5.8% of the heterozygote loci that passed the minimum filtration criteria in each individual (0.1% of the total expressed). This portion of the transcriptomic variants belonged to an average of 103 genes in each tissue transcriptome (approx.1%) or 300 in each individual (Supplementary Fig.S5 and S6). As shown in Supplementary Fig.S6. expression level varies across tissues but does not affect the distribution of ASE SNVs.

**Figure 3.**
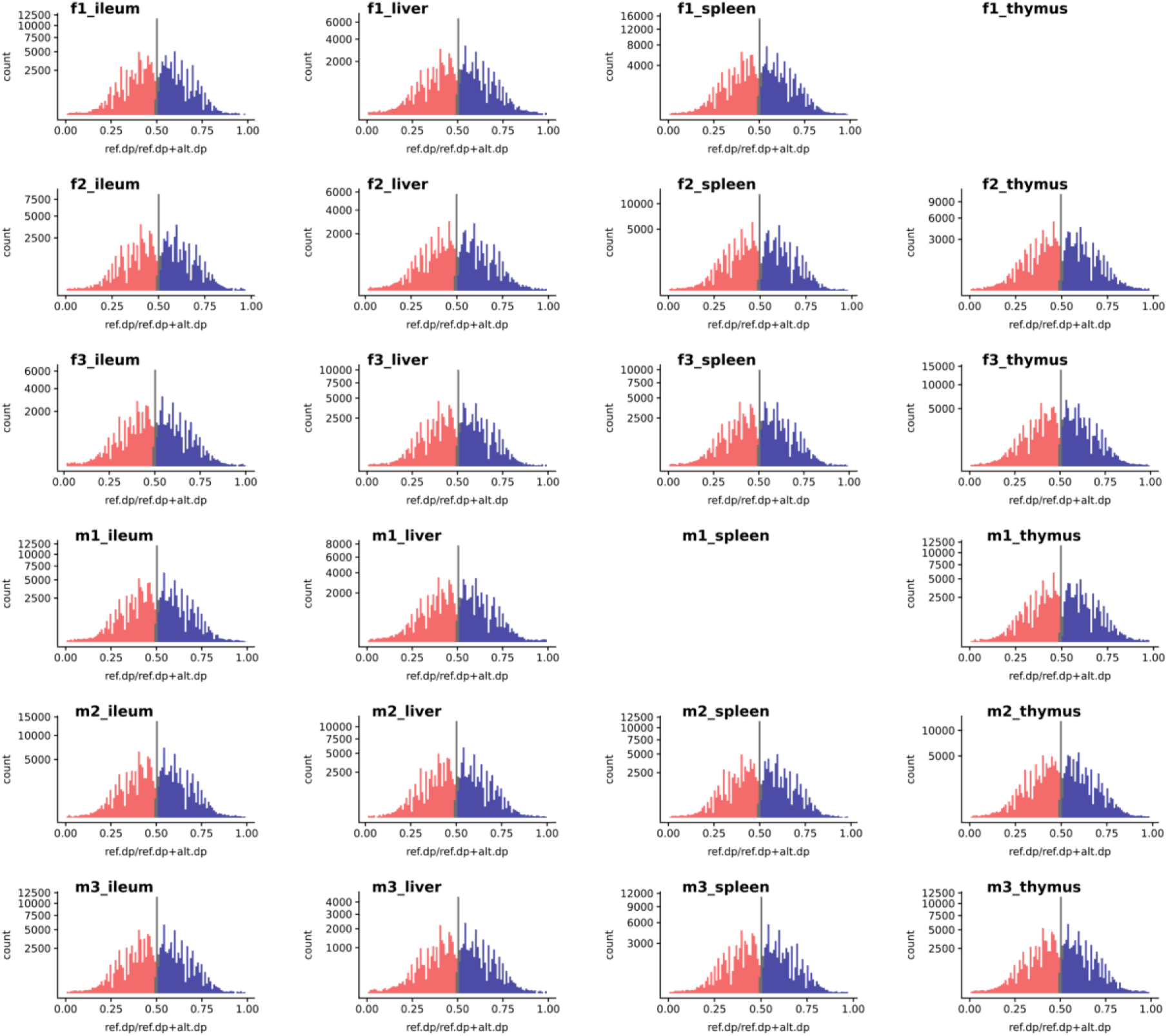
The histogram of a global reference allelic ratio at every locus in the tissues. The distribution of ref allelic ratio showed a balanced profile without any 0 or 1 inflation which is observed in the presence of reference mapping bias. The allelic ratio above 0.51 is shown in blue and below 0.49 in red while balanced bi-allelic expression (0.49-0.51) is coloured in grey. Ref.dp = read counts for reference allele. Alt.dp = read counts for alternate allele. The y axis is square root scaled. As discussed in the text SNP that display MAE are not present in any of the samples analysed, indicating there was no inflation in either 0 or 1 allelic ratio.

### Genes exhibiting tissue-specific and pervasive ASE signatures

We used the static mode of GeneiASE to investigate pervasive and tissue-specific ASE profiles across all of the available samples. Static ASE represents inherent allelic imbalance (AI) in each gene calculated by ASE at all heterozygote loci. The number of genes showing significant static ASE in immune-related tissues across the six sheep are summarised in (Table 1. On average approximately 0.5% of the genes in each tissue-specific transcriptome showed significant ASE (approx. 1% of the filtered set of genes). Pervasive ASE genes were investigated by applying the minimum 67% shared rule (i.e. a ASE gene was considered ‘shared’ when it exhibited ASE in a minimum of 4 out of 6 sheep). A list of ASE genes with significant allelic imbalance (AI) in all tissues, when the effect size was averaged across 6 sheep, was compiled (Fig.4A) (Static ASE measured by GeneiASE’s Liptak-Stouffer method). Six genes exhibited pervasive ASE across tissues (i.e. they were shared across all 4 tissues). In the order of allelic imbalance effect size they were NAA50 (N(alpha)-acetyltransferase 50, Histone acetyl transferase) with highest ASE effect size in spleen, UBB (Ubiquitin B, protein degradation) in thymus, HBP1 (HMG-Box transcription repressor; cell cycle regulation) and ENSOARG00000016510 both in spleen, C1orf105 (Chromosome 1 Open Reading Frame 105) in ileum and MTIF2 (mitochondrial translational initiation factor 2) in thymus.

**Figure 4.**
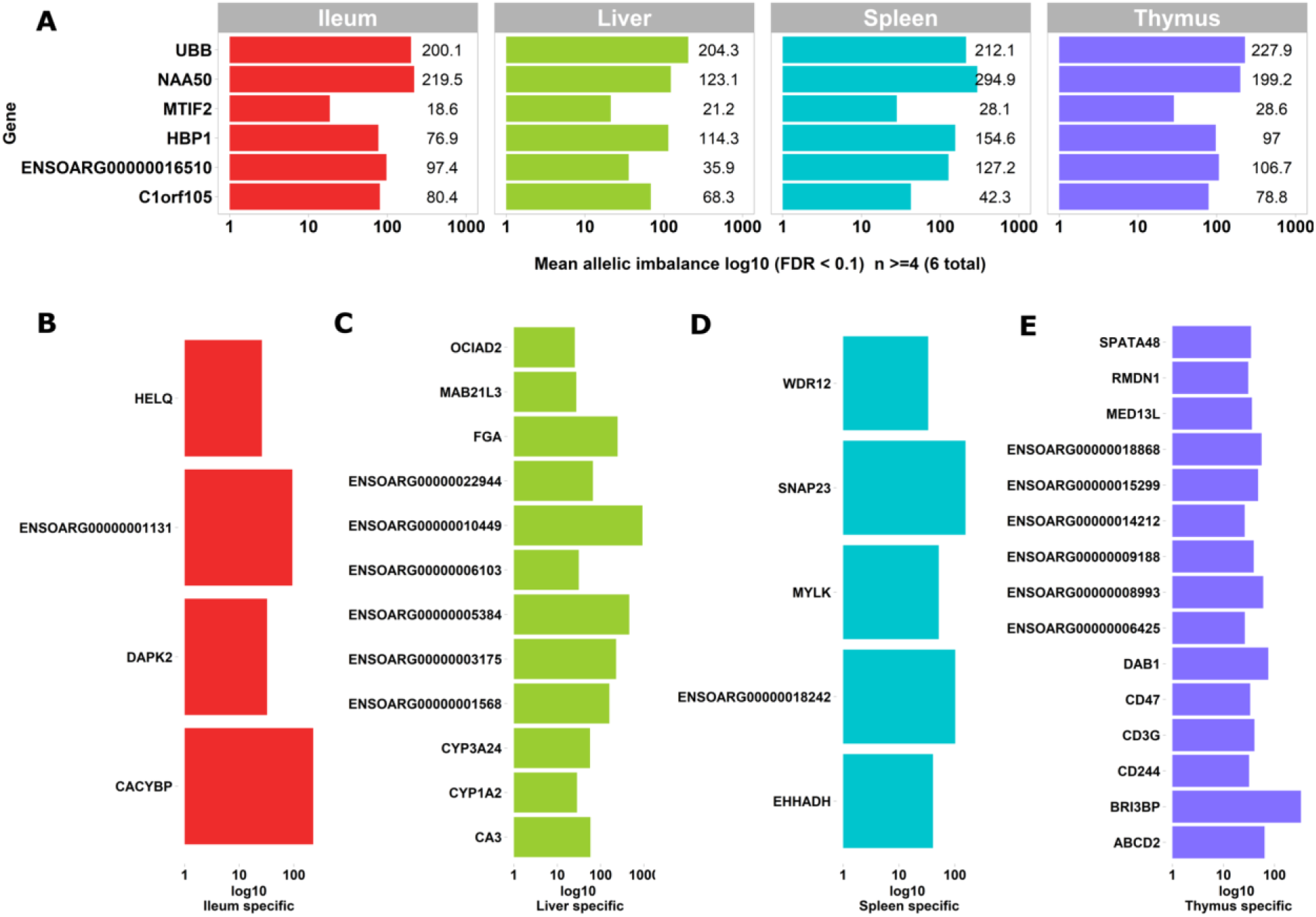
Genes exhibiting static ASE shared across tissues from all six sheep. The x axis represents the mean allelic imbalance (averaged static ASE across sheep in each tissue). **A)** genes shared by four tissues with significant (FDR < 0.1) static ASE. **B)** ASE genes private to Ileum. **C)** Private to liver. **D)** Private to spleen. **E)** Private to thymus.

**Tablel 1.**
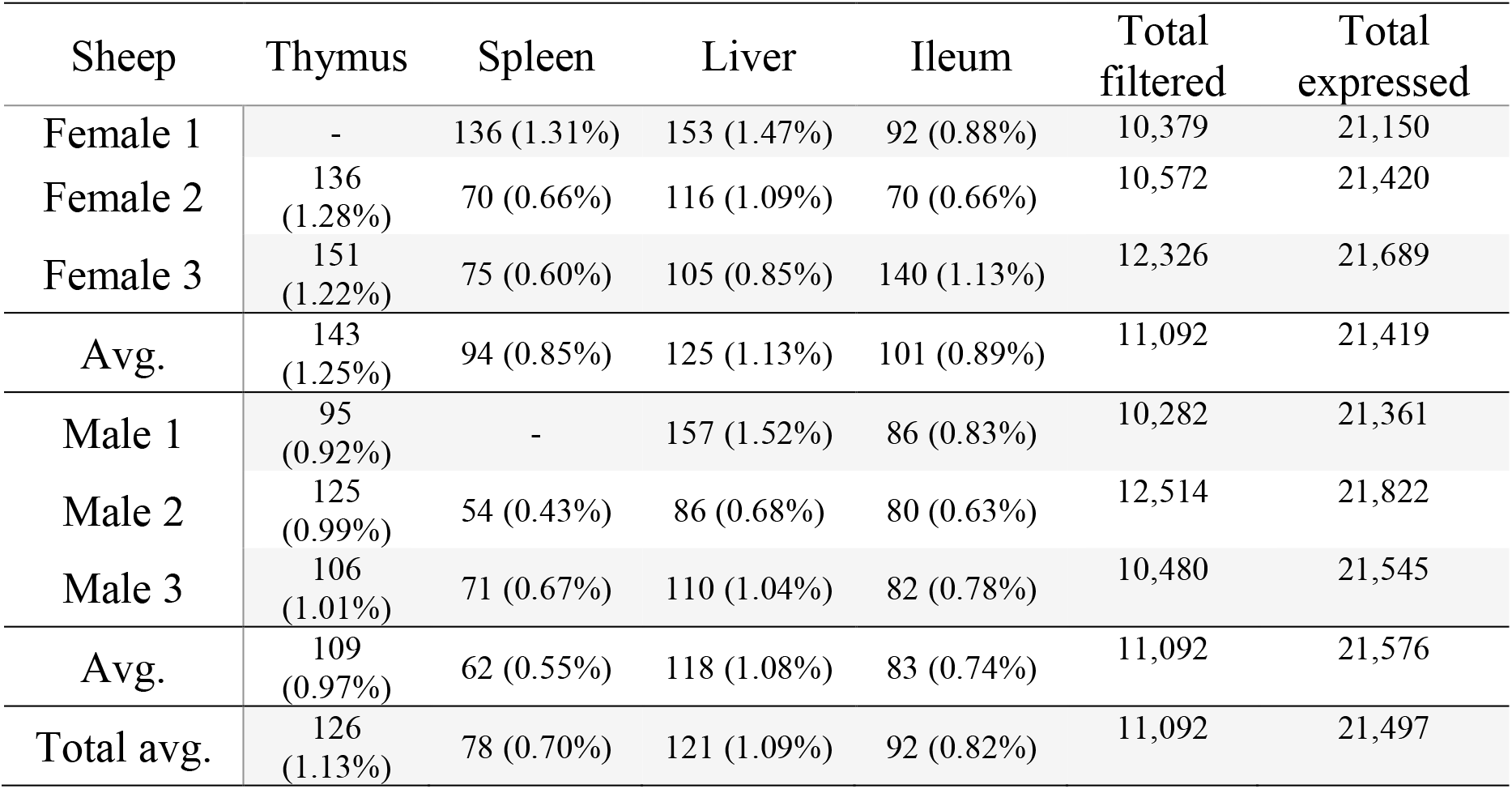
Total number of genes with significant static ASE in proportion to genes containing informative SNVs (filtered). Total expressed: Average number of genes being expressed in all 4 tissues. Total filtered: Average number of genes (containing heterozygote loci) passing read bi-allelic filtration criteria in 4 tissues. Tissue break-down has been presented as count (%ASE/filtered).

Sets of genes with tissue-specific ASE profiles were also captured (Fig.4B-E). Thymus had the highest number of tissue-specific ASE genes (n=15) followed by liver (n=12), spleen (n=5) and ileum (n=4) (Fig.4B-E). Amongst the thymus gene set was CD244 which included 30 heterozygote loci with allelic imbalance, one of which was rs406633825. This missense allele (Chr1:110308273 C>A; pVal123Phe MAF=0.7, SIFT score=0 deleterious) has previously been reported in the Texel population characterised by the International Sheep Genome Consortium (ISGC) (Kijas et al., 2012). The CD244, a non-MHC (major histocompatibility complex) mediated marker expressed by NK cells and multiple subsets of CD8+T cells is known for both pro-inflammatory and inhibitory effects on lymphocytes (McNerney et al., 2005; Georgoudaki et al., 2015). CD244 exons 2 to 5 are highly conserved in vertebrates and in mouse a Trypanosoma infection model indicated differential expression was correlated with multiple copy number variants nearby (Goodhead et al., 2010).The liver-specific ASE profile included genes involved in amino acid metabolism, cytochrome oxidase pathways and fibrinogen: FGA (fibrinogen alpha chain), ENSOARG00000003175 (taurochenodeoxycholic 6 alpha-hydroxylase-like, cytochrome P450 family 4 member [CYP4]), ENSOARG00000001568 (novel gene, complement C4-A-like), CYP3A24 (cytochrome P450 family 3 member) and CA3 (carbonic anhydrase 3). Allelic imbalance in spleen was present in CACYBP (calcyclin binding protein), DAPK2 (death associated protein kinase [serine/threonine]) and a novel gene GIMAP8-like (ENSOARG0000001131). The ASE in GIMAP8 has been previously reported in cattle with a strong paternal parent-of-origin expression pattern (Chamberlain et al., 2015). The GIMAP/IAN family of proteins, are involved in survival, selection and homeostasis of lymphocytes (Nitta and Takahama, 2007).

Two genes of functional interest showed evidence of strong tissue-specific ASE in the spleen: SNAP23 and MYLK. SNAP23 is a key molecule in vesicle transport machinery of the cell and has been reported to be expressed in sheep spleen. SNAP23 or Synaptosome Associated Protein 23 is part of protein complex involved in Class 1 MHC mediated antigen processing and presentation and in neutrophil degranulation (Fabregat et al., 2018). SNAP23 is also vital to lymphocyte development (both B and T) in vitro (Wong et al., 1997; Kaul et al., 2015). The myosin light chain kinase (MYLK) expression in the splenic trabeculae’s smooth muscle has been demonstrated previously (Jiang et al., 2014; Clark et al., 2017). Overall 199 heterozygote bi-allelic loci were present within the MYLK gene. The variant rs400678033 (Chr1:186347056G>A;.pAla1014Val), a missense SNV in exon 17 of 33 exons in MYLK, showed consistent allelic imbalance in all spleen samples.

In summary, analysis of ASE across immune related tissues revealed there were a small number of genes that were shared across tissues. ASE signatures instead tended to be tissue-specific, within the sub-set of tissues investigated in this study.

### Individual-specific ASE signatures

In order to investigate whether ASE profiles were either shared across all six sheep or private to individual sheep we used intersectionality (Fig.5). Each tissue was investigated separately. A number of private (to each individual) ASE genes were detected for each tissue, ranging from: 31-123 in ileum (Fig.5A), 24-80 in liver (Fig.5B), 21-83 in spleen (Fig.5C) and 31-66 in thymus (Fig.5D). Some of the shared sets of ASE genes in these tissues were specific to either male or female sheep, these sex-specific ASE signatures are described in Fig.5. In ileum no sex-specific set was observed (Fig5A). In contrast to ileum, the ASE profile for liver included a single gene with female only membership, ENSOARG00000017409 (novel gene; 93% orthology with bovine dicarbonyl and L-xylulose reductase [DCXR]) Fig.5B. In spleen all female sheep shared significant ASE in PMS1 (DNA mismatch repair system component during post meiotic segregation), ANKRD10 (Ankyrin Repeat Domain 10) and ENSOARG00000006103 which was not present in any of the spleen profiles of male sheep (Fig.5C). In thymus there were 2 sex-specific sets: 16 genes showing ASE only in females and 5 genes only in males. The female-specific thymus gene set included: ARPP21 (cAMP regulated phosphoprotein 21), CDKL3 (cyclin dependent kinase like 3), CEP19, ENSOARG00000000710 (novel gene), ENSOARG00000001163 (novel gene), ENSOARG00000000710, ENSOARG00000001163, ENSOARG00000008981 (novel gene; T-lymphocyte surface antigen Ly-9-like), ENSOARG00000006215, ENA000000008981, ENSOARG00000009129, ENSOARG00000011375 (blood vessel epicardial substance [BVES]), ENSOARG00000015755, ENSOARG00000020354 (novel gene; 53% orthology with bovine monoacylglycerol acyltransferase [MOGAT1]), ENSOARG00000025005, ENSOARG00000026030 (novel gene), GPM6A (glycoprotein M6A), RAG1 (recombination Activating 1), STX8 (syntaxin 8). The male-specific thymus set was comprised of ENSOARG00000007267 (novel gene; T-cell surface glycoprotein CD1a-like), ENSOARG00000016841 (novel gene; 98% orthology with bovine ATP synthase membrane subunit G [ATP5MG]), ENSOARG00000007603, SNX25 (sorting nexin 25) and LDHA (lactate dehydrogenase A) Fig.5D.

**Figure 5.**
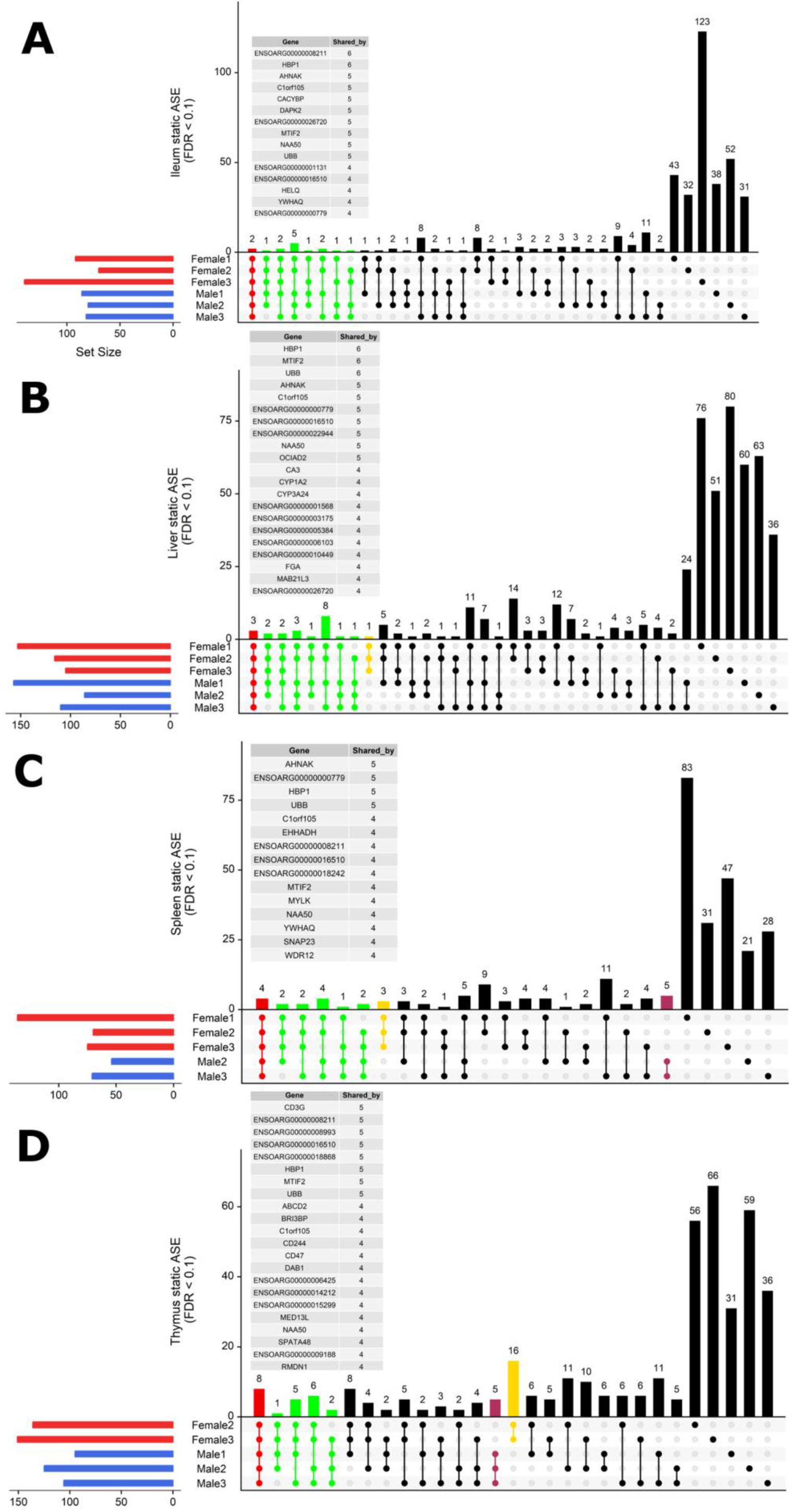
Intersectionality analysis of genes expressing significant ASE across all six sheep. In each tissue from left to right the set count of genes (dots connected by lines) illustrates the number of sheep sharing the gene. The private sets of genes are located at the far right of each graph (single dots with no line). The intersections are coloured in to illustrate the size of the set of shared genes (red [common to all 6 sheep], green [shared by 5 or 4 sheep], yellow [only in females] and purple [only in males]). Detailed lists of genes with ASE shared by at least 4 sheep are presented above each graph for **A)** Ileum, **B)** Liver, **C)** Spleen and **D)** Thymus. Two sex-specific sets of genes are highlighted: 16 genes showing ASE only in females (in yellow) and 5 genes only in males (in purple).

In summary very few ASE genes were shared across all sheep and the majority of ASE profiles were private to each sheep. Sex-specific ASE signatures were also detected but due to the small sample size (n=3) in both cases these should be interpreted with caution.

### ASE in stimulated and unstimulated BMDMs (0hrs vs 7hrs +LPS)

We examined inducible ASE after 7 hrs of exposure to LPS in BMDMs using the ICD mode of GeneiASE (Edsgärd et al., 2016). A comparison of LPS inducible ICD-ASE genes and the genes with background static ASE at 0 and 7 hr time points, was also performed. We first assessed whether differences between 0 and 7 hours could be observed using analysis of static ASE. Individual-specific ASE profiles and a limited number of shared ASE genes were also observed in BMDMs. The total number of genes with static ASE in the BMDMs is shown in (Table 2.

**Tablel 2.**
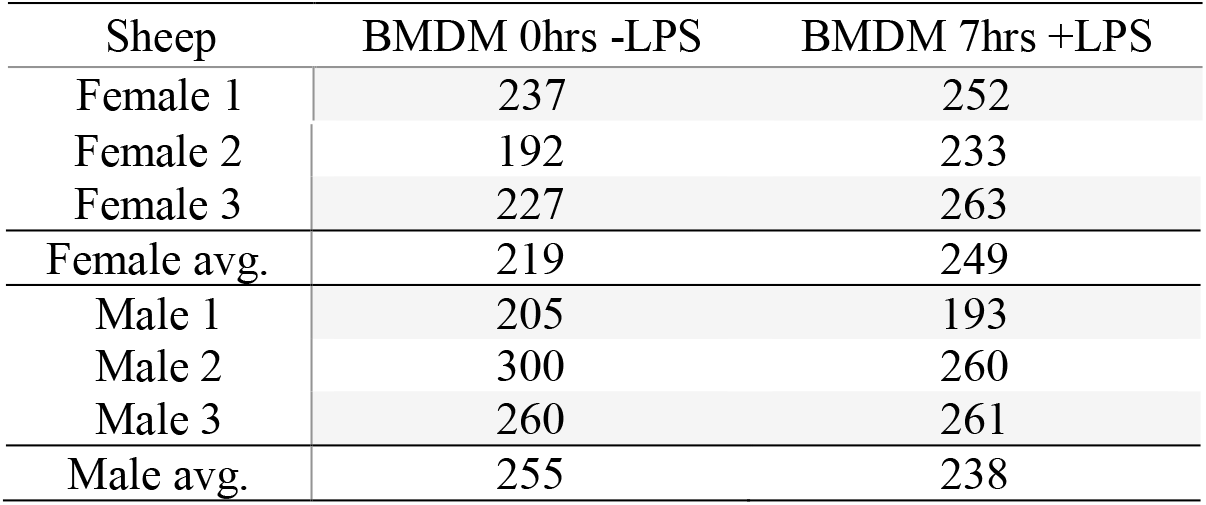
Total number of genes with significant static ASE in BMDMs +/− LPS.

Shared static ASE across both time points and independent of LPS induction was only observed in 5 genes. These genes have a macrophage associated function and include ITGB2 (Yee and Hamerman, 2013), SAA3 (ENSOARG00000009963) (Larson et al., 2003; Deguchi et al., 2013), CD200R1 (ENSOARG00000019357) (Ocaña-Guzman et al., 2018), DCTN5 (ENSOARG00000017281) (Habermann et al., 2001) and MTIF2 (also seen in the tissue analysis above) (Overman et al., 2003).

The ICD-ASE in BMDMs +/−LPS captured fewer ASE genes with significant LPS inducible ASE between the 2 timepoints than the static analysis of ASE. Moreover, there were large differences in the number of LPS inducible ASE genes in each individual sheep, indicating significant individual-specific variation in response to LPS stimulation. BMDM cultured from female 2 showed no LPS inducible response in comparison to male 3 which was a hyper responder with significant inducible ICD-ASE in 28 genes (including 634 informative SNVs total). A detailed breakdown of SNVs, aggregated within each gene, with significant ICD ASE has been summarised in Fig.6.

**Figure 6.**
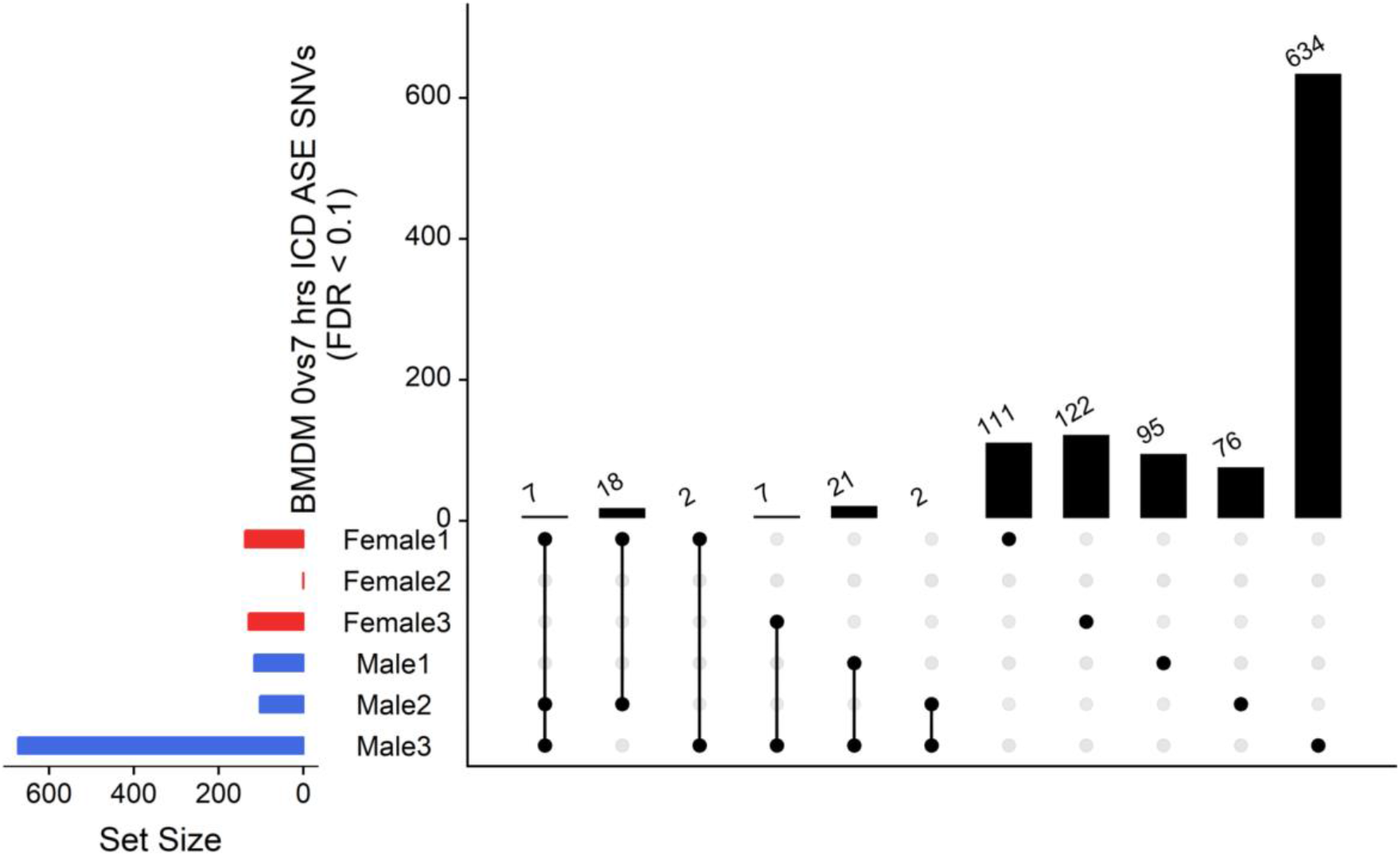
Intersection analysis of SNVs under genes with significant ICD-ASE in the BMDMs ± LPS. From left to right the set number of genes (dots connected by lines) have been illustrated in order according to the number of sheep sharing the SNV. The private sets of SNVs are located at the far right of each graph (single dots with no line).

In summary the ICD-ASE mode of GeneiASE’s model was not capable of capturing a complete picture of differential ASE in the BMDM experiment. Static ASE was present in both time points, however, there were no inducible ASE genes that were shared across all 6 sheep. The highly diverse ASE profile of BMDMs was very individual-specific, similar to patterns observed in tissues (Supplementary Table S1). These individual-specific differences could be due to individual variation in the innate immune response or experimental variation introduced during primary cell culture or stimulation with LPS.

### Condition dependent ASE at the SNV level in BMDMs

To further investigate allelic imbalance at the SNV level without the aggregative gene model of GeneiASE, Fisher’s exact test was used. The filtered read counts for bi-allelic SNVs shared by all 6 sheep BMDMs were selected (n=646 sites). Allelic read counts of each SNV were tested using Fisher’s exact test between 0hrs and 7hrs (2×2 table). Overall the 6 sheep shared 646 SNVs with identical allelic genotypes in both time-points of the BMDM RNA-Seq dataset. These SNVs were tested for association with LPS treatment and only 4 SNVs showed a strong association (FDR < 1.0e-8) and 12 SNVs had an FDR between 1.0e-02 and 1.0e-08 (Fig.7). The highest F-statistic was at rs430667535 Chr17 Pos:50485358T>C, a synonymous variant in ubiquitin C (UBC), a polyubiquitin precursor, and also an intronic variant T>C or A>G in pro-apoptotic BRI3 binding protein (BRI3BP). This variant was shown to have a minor allele frequency (MAF) of 0.25 in Texel sheep based on the ISCG annotation [ISGC – Ensembl v.92](Kijas et al., 2012)). The next highest peak was observed on Chr21 under SNVs within SAA3 gene boundaries (ENSOARG00000009963) at the following coordinates: Highest FDR peak was observed at SNV rs412192652 (Pos:25826978A>G missense variant [Asn145Asp], FDR=2.3e-15) surrounded by rs403064928 (Pos:25826884C>A missense variant [Asp113Glu] in exon 4 of SAA3 FDR=3.6e-07), rs426609498 (Pos:25826845A>G synonymous, FDR=5.5e-07) and rs405439099 (Pos:25826990G>C 3’UTR variant, MAF 0.6 in Texel sheep, FDR=4.1e-05). This region contains a strong LD block previously reported by the ISGC COMPOSITE population (Ensembl v.92) (Supplementary Fig.S4) e.g. rs412192652 and rs405439099 pairwise D’ statistics = 1. Two further peaks on Chr3 were observed for rs159926581 (Pos:214731375T>C synonymous, FDR = 3.3e-09) in ribosomal protein L3 (ENSOARG00000016495) and rs159822214 (Pos:112164732G>A 3’UTR variant, FDR=9.5e-03) in oxysterol binding protein like 8 (OSBPL8). The last inducible ASE associated signal was on Chr16 rs420037698 (Pos:6887423G >A, FDR= 6.9E-09) in ENSOARG00000004700, a known synonymous variant in the Texel population (MAF = 0.55). The SNVs and corresponding genes from Fisher’s exact test are summarized in Table 3.

**Figure 7.**
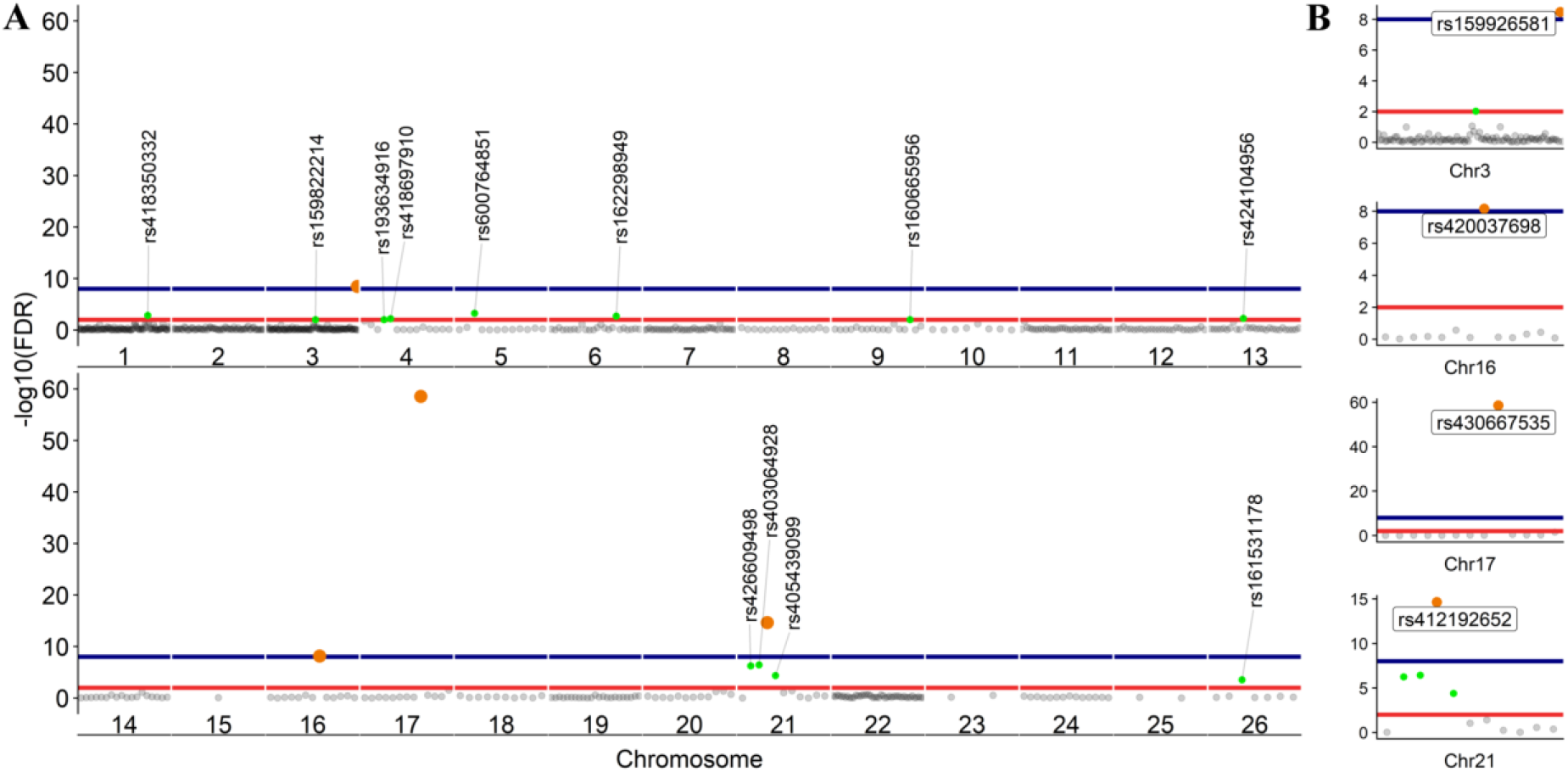
Scatter plot of the adjusted p values from Fisher’s exact test (unified using Stouffer unification) in BMDMs comparing expression from different alleles at 0 vs 7hrs at SNV level (LPS inducible ASE). **A)** The graph shows 646 loci exhibiting LPS inducible allelic imbalance shared across all 6 sheep. **B)** Four loci on chromosomes 3,16,17 and 21 with FDR < 1.0e-08. FDR < 1.0e-02 red line (n=16 SNVs) and FDR < 1.0e-08 blue line (n = 4 SNVs).

**Tablel 3.**
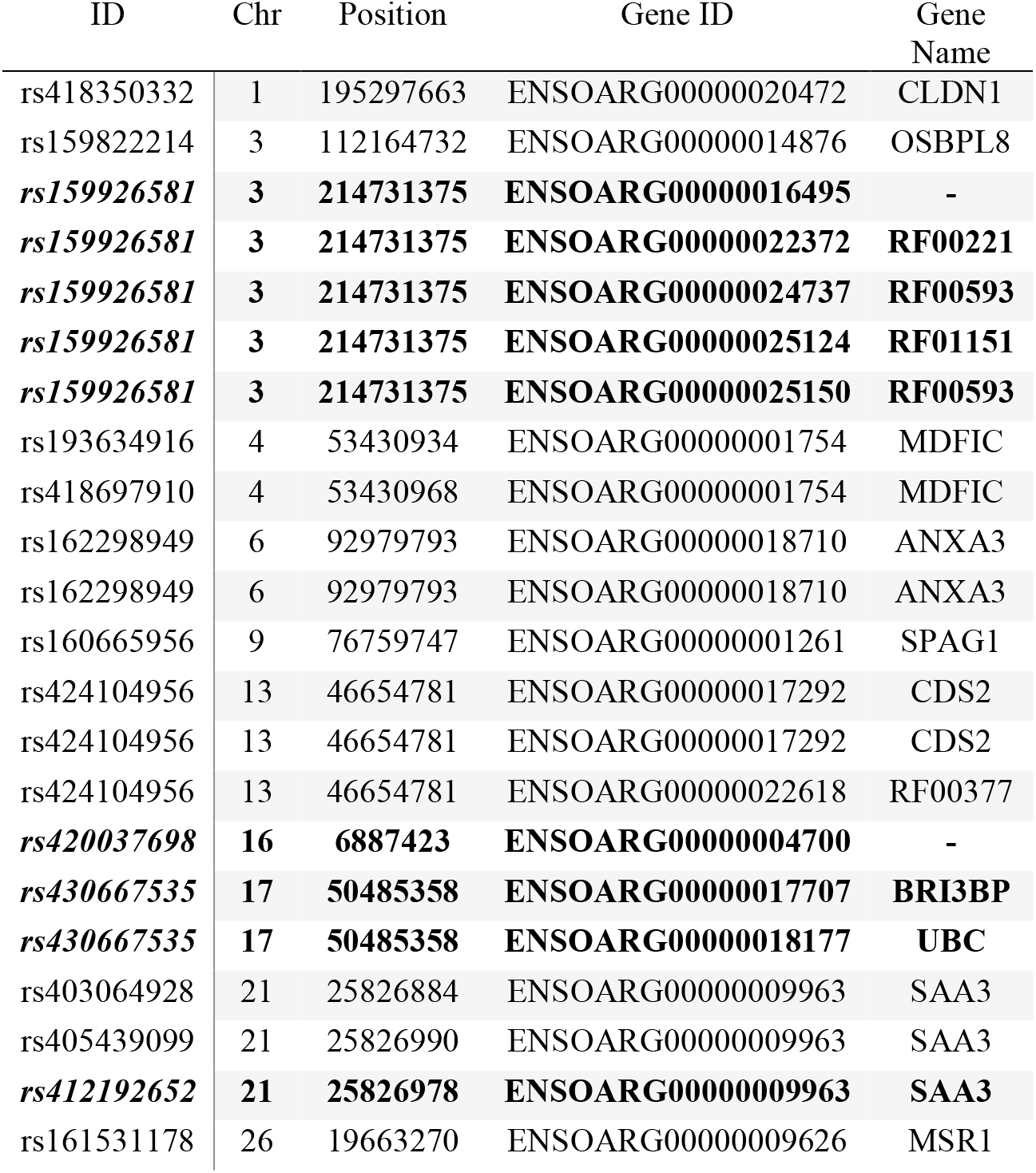
The variant IDs of LPS inducible ASE SNVs (Fisher’s exact) and their respective genes. Data was obtained using Ensembl BioMart query builder. Highly significant SNVs are highlighted in bold (FDR< 1.0e-08)

The LPS inducible ASE analysis, of SNVs, using Fisher’s exact test revealed a different picture not captured by the gene level analysis with the GeneiASE model that aggregates SNVs within each gene (Fig.6 vs Fig.7). The aggregative gene model did not capture any shared ASE genes in the ICD-ASE mode. Although the 6 sheep shared 646 SNVs and showed highly significant association with stimulation with the LPS (Fisher’s exact method), the aggregation of ASE effect size from SNV to gene level (ICD-ASE mode) only detected individual-specific sets of ASE genes in each sheep. This contradicted the results from Fisher’s exact test which detected 4 highly significant LPS inducible shared regions (FDR < 0.01, 16 SNVs) on chromosomes 3, 16, 17 and 21. For example, the allelic imbalance in the SAA3 genomic coordinates on chromosome 21 was not detectable in the ICD-ASE model but it was captured by the Fisher’s exact test in all individuals (Fig.7B chromosome 21).

Fisher’s exact analysis at SNV level revealed ASE in response to LPS in variants within CLDN1, ANXA3, BRI3BP, SAA3 and MSR1 (Table3). The anti-inflammatory macrophage marker CLDN1 (Van den Bossche et al., 2012) and acute-phase inflammation resolution marker ANXA3 (Yamanegi et al., 2018) have been previously reported with distinct macrophage functions. The macrophage scavenger receptor 1 (MSR1) has also been shown to be involved in lipid uptake and migration ability of macrophages (Shi et al., 2019). Three noncoding RNAs (RF00221 [snoRD43], RF00593[snoU83B] and RF01151[snoU82P] were amongst genes corresponding to ASE inducible SNVs. These 3 snoRNAs all overlap with the genomic coordinates of ribosomal protein L3 (ENSOARG00000016495 / RPL3). The RF00377 [snoU6-53] was also among the ASE positive targets which overlaps the protein-coding gene CDS2 (CDP-diacylglycerol synthase 2). Using total RNA-Seq (ribosomal RNA depleted), which includes multiple RNA populations, to generate short read illumina data makes it difficult to pinpoint the origin of the ASE signal to a specific RNA population.

In summary ASE profiles in BMDMs were highly individual-specific at both gene and SNV level. Moreover, Fisher’s exact SNV level analysis discovered shared ASE SNVs where the aggregative gene model of ICD-ASE mode did not, indicating for condition dependant ASE analysis Fisher’s exact test is more accurate and robust at SNV level.

## Discussion

This study is the first to investigate global allele-specific expression across tissues from sheep using RNA-Seq data. We focused our analysis on immune-related tissues and cell types from six adult crossbred sheep (TxBF) from the sheep gene expression atlas. ASE profiles were highly individual-specific in the six sheep analysed in this study. We were able to identify tissue-specific sets of ASE genes, as well as LPS inducible sets in the BMDM experiment. Tissue-specific signatures of ASE have been previously reported in similar studies in mouse (Castel et al., 2015, 2016), goat (Cao et al., 2019) and cattle (Chamberlain et al., 2015).

Several steps were taken in the cattle study (Chamberlain et al., 2015) to mitigate the ref allele bias, assign parental origin using whole genome sequences and include MAE variants. The SNV filtration was based on the (Hayes and Daetwyler, 2019) 1000 bull genomes project to confirm the heterozygote sites. In our pipeline the Ensembl VCF track was used for that purpose. Chamberlain et al., 2015 use a 0.9 allele frequency cut-off (based on read counts) to define and include MAE, and as such have a 1 and 0 inflated allelic ratios. In our pipeline no allelic ratio cut-off is introduced for inclusion as it is difficult to distinguish between sequencing error and MAE. The minimum read (bi-allelic expression) filtration criteria was applied to exclude highly sequenced loci (Either count/Total >1%) or sequencing errors presenting as rare alleles (min either allele count >=3) which consequentially excludes actual MAE as well as spurious allelic counts. Chamberlain et al. tested 5317 genes (14,495 SNVs) in spleen and detected 382 ASE genes (with min > 1 SNV per gene, similar to this study). Although direct comparison would not be appropriate (because we have excluded MAE variants in our analysis) our analysis of sheep spleen revealed ASE in 86 genes (averaged in 5 sheep) from 8272 filtered genes (averaged in 5 sheep). Similarly, in thymus the cattle study showed 182 ASE genes from 986 informative genes (9781 SNVs) while from 7961 filtered genes in sheep thymus 134 ASE genes were captured. The differences in the numbers of genes exhibiting ASE between the two studies are likely to be a consequence of the filtration criteria applied, the exclusion of MAE, and species-specific differences between sheep and cow. Results from a more recent study in goat (Cao et al., 2019) more closely reflect our findings. They apply similar filtration criteria to our workflow and discovered 144 ASE genes in liver in comparison to 123 in our sheep liver sets (averaged across 6 sheep). Other recent studies including those focusing on production relevant tissues, such as muscle (Guillocheau et al., 2019), have also applied similar stringency in filtration criteria. The filtering criteria we have used for this analysis is stringent and focused on detecting variants of moderate to extreme effects. Further analysis of the dataset reducing these criteria might discover additional variants exhibiting ASE across individuals and tissues, but it would also increase the potential risk of false positive discovery.

For this analysis we have adapted an ASE analysis workflow with a primary focus on mapping bias removal prior to allele-specific analysis of the transcriptome. The collection of scripts for WASP, used for this analysis, or modified versions of them have been utilised by others for mapping bias removal in reference guided genomic datasets e.g. RNA-Seq (Mozaffari et al., 2018; Zhou et al., 2018), Chip-Seq (Pelikan et al., 2018) and for methylomic and epigenetic analysis (Richard Albert et al., 2018).

The ASE analysis pipeline we have adapted for sheep for this study is also adaptable to other species and tissue types with available RNA-Seq datasets. It could be applied, for example, to profile allele-specific expression in the RNA-Seq datasets from livestock species listed on the FAANG data portal (Andersson et al., 2015; Harrison et al., 2018). We used the Ensembl VCF track to capture information at heterozygote loci however the individual VCF file from each sheep could also be used in ASE analysis. The latter strategy might enable the capture of rare variants not included in the publicly available VCF tracks but would also raise the issue of normalisation/standardization between VCF call sets. The usage of either of these methods will be limited to the number of loci shared by coordinate and bi-allelic genotype (i.e. pervasive ASE discovery). Other studies have compared variants at the RNA and DNA level from the same individual and then removed the DNA variants not present in the RNA-Seq data from the analysis (Guillocheau et al., 2019). We believe that a strength of the pipeline we present is that it does not require parental genotypes and can therefore be applied to other RNA-Seq datasets for livestock where this information is not available.

In our analysis we have not considered either parent-of-origin or breed-of-origin-specific effects in this analysis. For parent-of-origin or breed-of-origin assignment of these ASE profiles DNA level genotypes from the parents of the six sheep from the gene expression atlas (i.e. Texel sire and Scottish Blackface dam) would be required, and these are unfortunately not available. In this study ASE expression profiles also might be affected by the direction of the cross (i.e. Texel sire x Scottish Blackface dam). To fully characterise parent-of-origin or breed-of-origin reciprocal cross experiments would be required. Reciprocal cross studies in mouse (Huang et al., 2017), chicken (Zhuo et al., 2017) and crossbred cattle (Chen et al., 2016b) have shed light on the complexity of such pervasive ASE markers and parent-of-origin effects. Though potentially very interesting these experiments are lengthy and costly to perform in sheep. Particularly in this case because the reciprocal cross (Scottish Blackface dam x Texel sire) is rarely used in the UK sheep industry and as a consequence has limited relevance to production.

Our approach also excludes mono-allelic expression. The minimum filtration criteria utilised in our workflow along with the reference mapping bias removal step ensures an unbiased ASE discovery in the transcriptome by excluding the ambiguity surrounding MAE variants. This form of analysis is based on the principle that absence of evidence (reads) for either allele of a heterozygote site does not directly amount to evidence of their absence i.e. MAE. The pattern of ASE (ratio of Alt/Ref+Alt) is dependent on the bi-allelic expression of loci within the genomic coordinates of the gene or genomic element of interest. For an ASE effect to be captured by the GeneiASE model, the following criteria must be met: i) Biallelic expression of the locus; ii) Min depth criteria for each allele (min 3 reads, total 10 reads at that site and > 1% of total reads containing that allele); iii) The allelic imbalance or departure from bi-allelic balanced expression being inducible by an environmental trigger (i.e. LPS in ICD ASE experiment with BMDM data). These stringent criteria secure robust transcriptome-wide ASE discovery while maximising the usage of read counts from short read RNA-Seq datasets without considering mono-allelic sites. MAE patterns are impossible to differentiate from sequencing error or random nonsense mediated decay in total RNA-Seq, unless arbitrary cut-offs are introduced such as ratio of allelic read counts > 0.9 (Chamberlain et al., 2015) or > 0.7 (Cao et al., 2019). We decided to exclude MAE from this study using read count bi-allelic expression filtration because it is difficult to distinguish between sequencing error and MAE. We do appreciate that this form of filtration might lead to a reduced number of ASE genes discoveries overall and will exclude potentially imprinted loci altogether.

The tissues utilised for ASE analysis in this study (thymus, spleen, liver and ileum) are highly influential on performance of the immune system. ASE profiles shared across tissues and cell types were limited and instead they tended to be highly specific. We identified tissue-specific ASE in several genes in Thymus, for example, that are involved in the T-cell mediated immune response, including CD47 and CD244. These tissue-specific and cell-type-specific ASE profiles may underlie the expression of economically important traits such as disease resistance. Assessment of the connection between economically relevant phenotypes and tissue-specific ASE profiles could be useful for the improvement of genomics enabled sheep breeding programmes, particularly those using specialised sire and dam lines (Georges et al., 2018). Loci exhibiting ASE have been associated with production traits including milk-fat percentage (Hayes et al., 2010; Suárez-Vega et al., 2017), trypanotolerance in small ruminants (Kadarmideen et al., 2011; Álvarez et al., 2016), mastitis in goat (Ilie et al., 2018), Johne’s disease in cattle (Mallikarjunappa et al., 2018) and Marek’s disease in chicken (Maceachern et al., 2011; Meydan et al., 2011; Cheng et al., 2015). Although there is no general consensus currently on the correlation of allelic expression haplotypes and phenotypes under selection in sheep, this form of ASE analysis could pave the way for functional validation at population level (e.g. breed or haplotype-specific aseQTL studies in a larger population of sheep). Examples of population level aseQTL, eQTL and sQTL (QTLs associated with RNA splicing) already exist for cattle (Wang et al., 2018; Xiang et al., 2018). Knowledge of favourable ASE in critical genes involved in traits of interest could be used as a performance indicator or included as weighted SNVs in genomic prediction algorithms to enhance livestock breeding programmes (Georges et al., 2018). Currently the UK sheep industry is on the cusp of applying genomic prediction, but suitable genomics enabled breeding programmes for sheep already exist in New Zealand and Australia (Daetwyler et al., 2010).

## Conclusions

In this study we characterise extreme to moderate allele-specific expression, at the gene and SNV level, in immune-related tissues and cells from six adult sheep (TxBF) from the sheep gene expression atlas dataset. Reference mapping bias removal was an integral component of the analysis pipeline applied in this study. The correction of reference bias prior to obtaining the allelic read counts is a critical step toward true ASE discovery. The workflow developed as part of this manuscript provides an RNA-Seq only dependent tool, without the need for individual DNA sequences. We note that the stringent filtering process applied would remove loci where the allelic imbalance was less extreme but might still be of biological significance.

This study is a novel analysis of an existing large-scale complex RNA-Seq dataset from sheep. Using the pipeline we have adapted for this analysis, we were able to identify ASE profiles that were pervasive in each sheep and specific to the tissues and cell types investigated. These tissue and cell-type-specific ASE profiles may underlie the expression of economically important traits and could be used to identify variants that could be weighted in genomic prediction algorithms for the improvement of sheep breeding programmes. In summary we have adapted a robust methodology for ASE profiling, using the sheep gene expression atlas dataset, and provided a foundation for identifying the regulatory and expressed elements of the genome that are driving complex traits in livestock.

## Data Availability

The RNA-Seq sequence data is available via the European Nucleotide Archive (ENA) under PRJEB19199 (https://www.ebi.ac.uk/ena/data/view/PRJEB19199). The BAM files were already produced as part of the sheep gene expression atlas following the publicly available protocol (FAANG, 2018) and as described in (Clark et al. 2017). The BAM files have been uploaded to ENA under accession numbers ERZ827944, ERZ827949, ERZ827951, ERZ827955, ERZ827972, ERZ827988, ERZ827995, ERZ827997, ERZ828001, ERZ828016, ERZ828019, ERZ828036, ERZ828044, ERZ828046, ERZ828050, ERZ828070, ERZ828073, ERZ828160, ERZ828167, ERZ828168, ERZ828172, ERZ828188, ERZ828192, ERZ828209, ERZ828215, ERZ828217, ERZ828221, ERZ828240, ERZ828244, ERZ828261, ERZ828268, ERZ828270, ERZ828274, ERZ828293 and ERZ828297. The Oar v3.1 reference FASTA and VCF file from Ensembl v92 were used throughout the pipeline. The ASE analysis pipeline (https://msalavat@bitbucket.org/msalavat/asewrap_public.git) was wrapped using bash scripting on Edinburgh Compute and Data Facility computing resource Eddie Mark 3 (Edinburgh, 2018). All the raw ASE genes data produced by GeneiASE could be found within Supplementary_file2.zip.

## Author’s contributions

MS and ELC coordinated and designed the analysis component of the study with assistance from SJB and SPV. MS and SPV designed, optimised and tested the ASE pipeline. DAH acquired the funding for the sheep gene expression atlas project. MEBM, ELC and DAH designed the LPS experiment and generated the data. MEBM performed the LPS stimulation of bone marrow derived macrophages and RNA extraction. SJB performed all bioinformatic analyses prior to analysis with the ASE pipeline. MS performed ASE analysis, visualisation of the results and wrote the manuscript. ELC contributed to manuscript editing and drafting. All authors read and approved the final manuscript.

## Acknowledgments

The authors would like to thank University of Edinburgh Computing and Data Support staff (viz. Dr Andy Law and Dr Steve Thorn) for their input towards optimisation of the pipeline and efficient parallel computing. Rachel Young and Lucas Lefevre isolated the bone marrow derived macrophages for the sheep gene expression atlas project and Iseabail Farquhar helped to collect and archive the tissue samples. The authors would also like to thank Professor Kim Summers and Professor Mick Watson for their comments, which helped to improve the manuscript.

## Supplementary material

A bimodal p value distribution was observed in GeneiASE output as shown in S1 Fig. 1:

**Figure S1.**
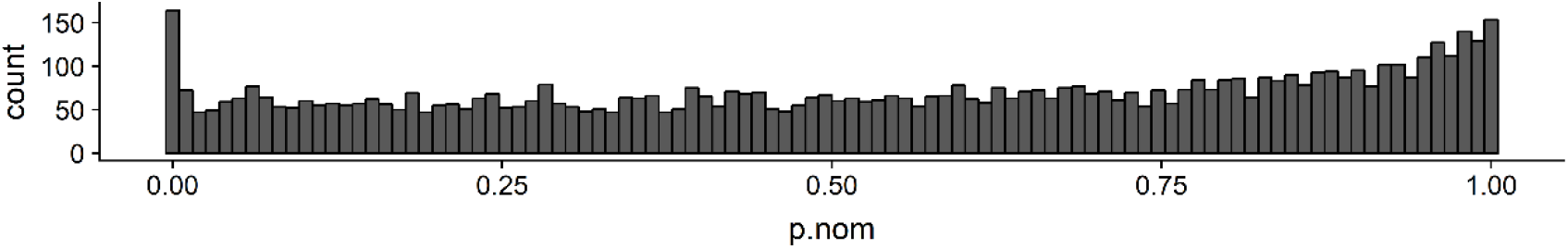
Distribution of p values resulted from the beta-binomial test (Static ASE example for spleen in female 1). The uniformity of the p value distribution showed a bimodal (high frequency close to 1 followed by an extensive lower tail close to 0). R machine characteristics for printing float numbers were limited to 6 significant numbers after the decimal point by default which resulted in a high number of absolute zero (0+e0.0) output-values.

**Figure S2.**
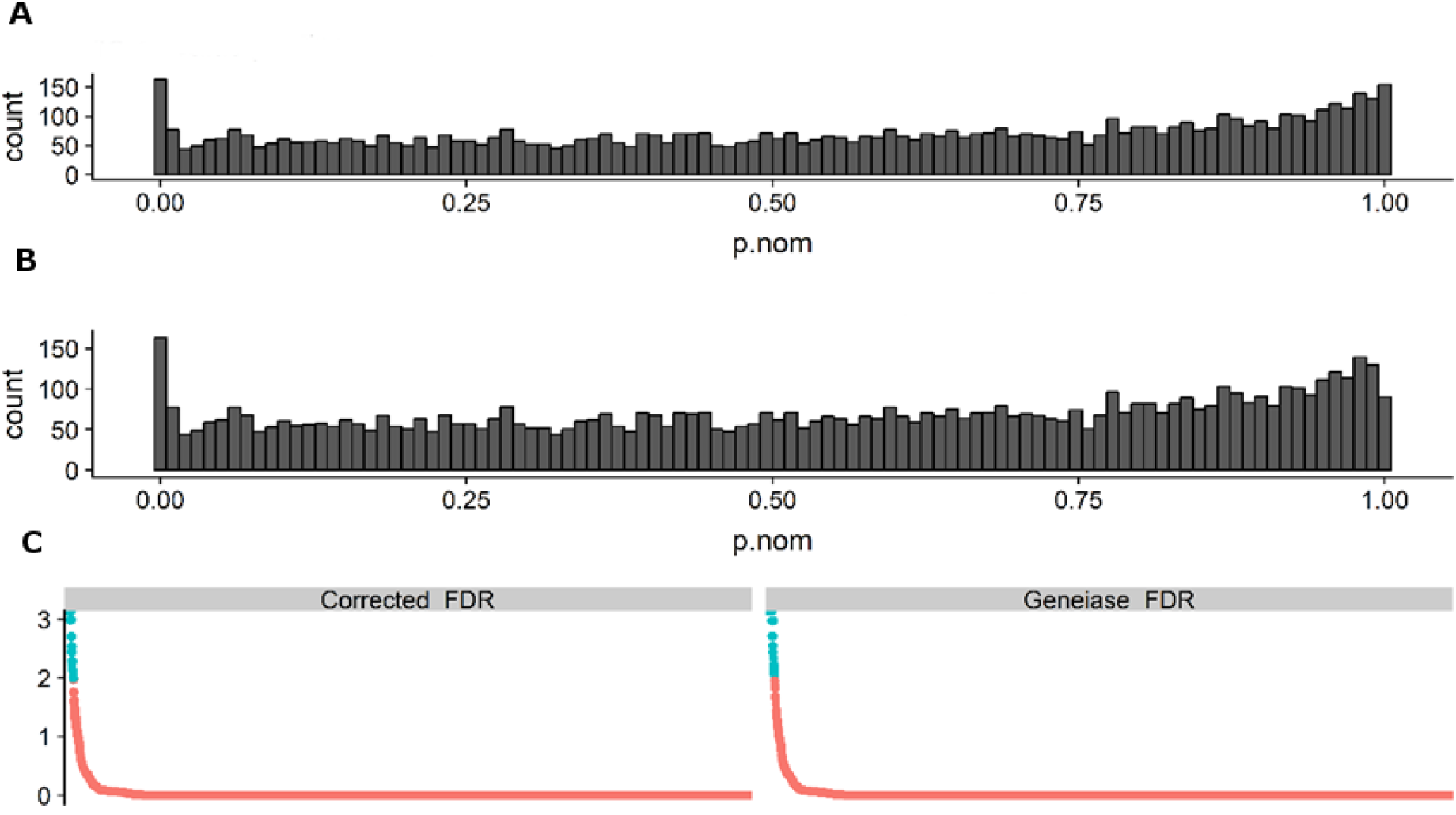
The figure shows the distribution of p values produced by GeneiASE after cut off introduction (A) and after removal of p value == 1 values (B). The distribution of sorted FDR calculated by GeneiASE and after correction are shown in C (right and left respectively). To correct for multiple testing by recalculation of FDR and eliminating the non-uniformity of p value distribution around 1 the following steps were performed as shown in S1 Fig.2: i) An arbitrary cut off (1e-7) was placed over all p values smaller than the said limit including the absolute 0e+0.0; ii) The genes showing p values equal to 1 were removed from each data frame; iii) The false discovery portion and rate were recalculated in each sample. iv) Gene showing FDR < 0.1 selected for further analysis.

**Figure S3.**
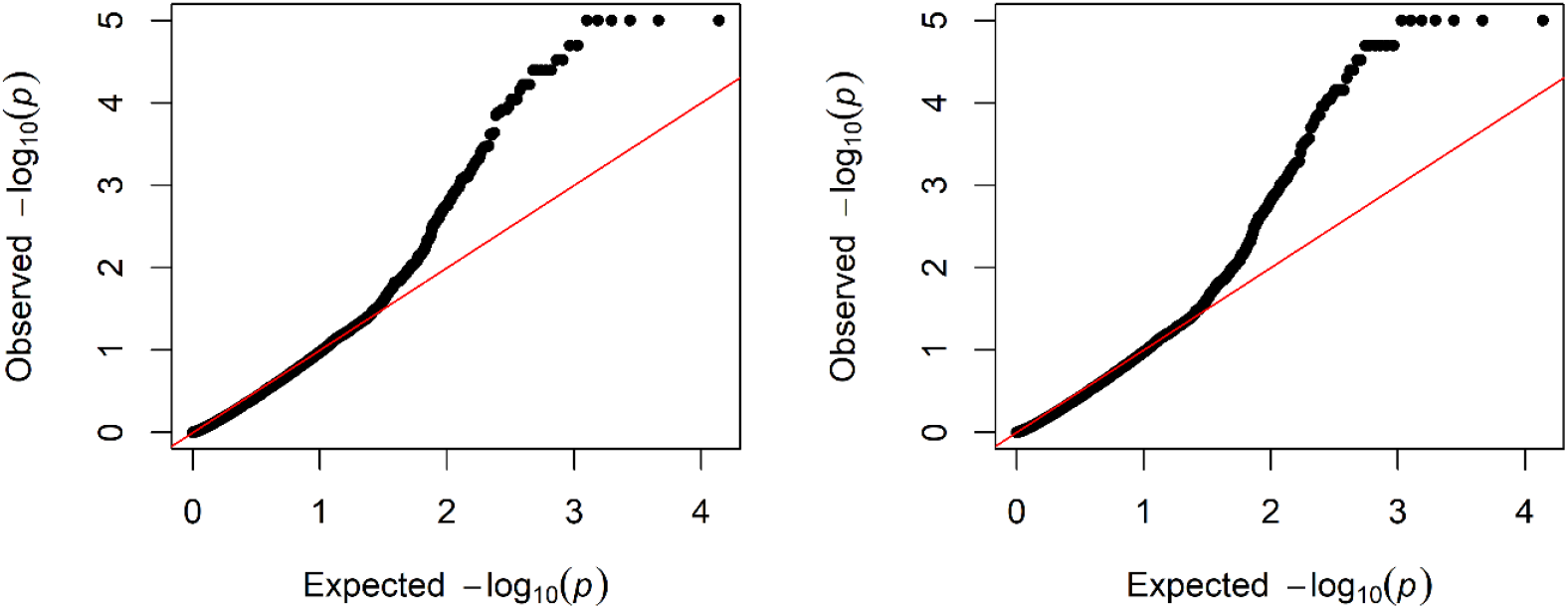
The QQ plot of observed vs expected p values in corrected (left) and GeneiASE’s raw (right) calculated outputs.

**Figure S4.**
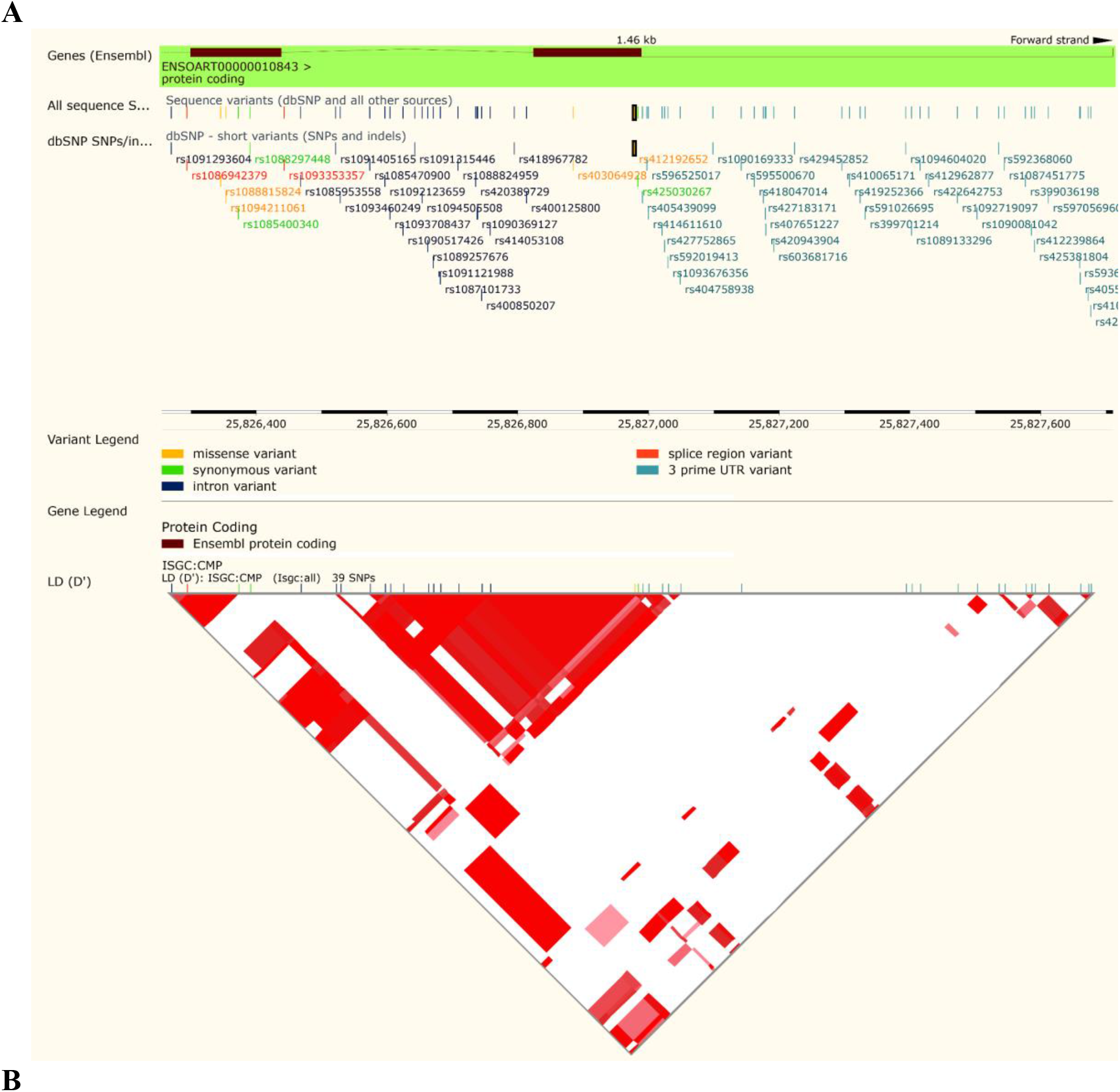
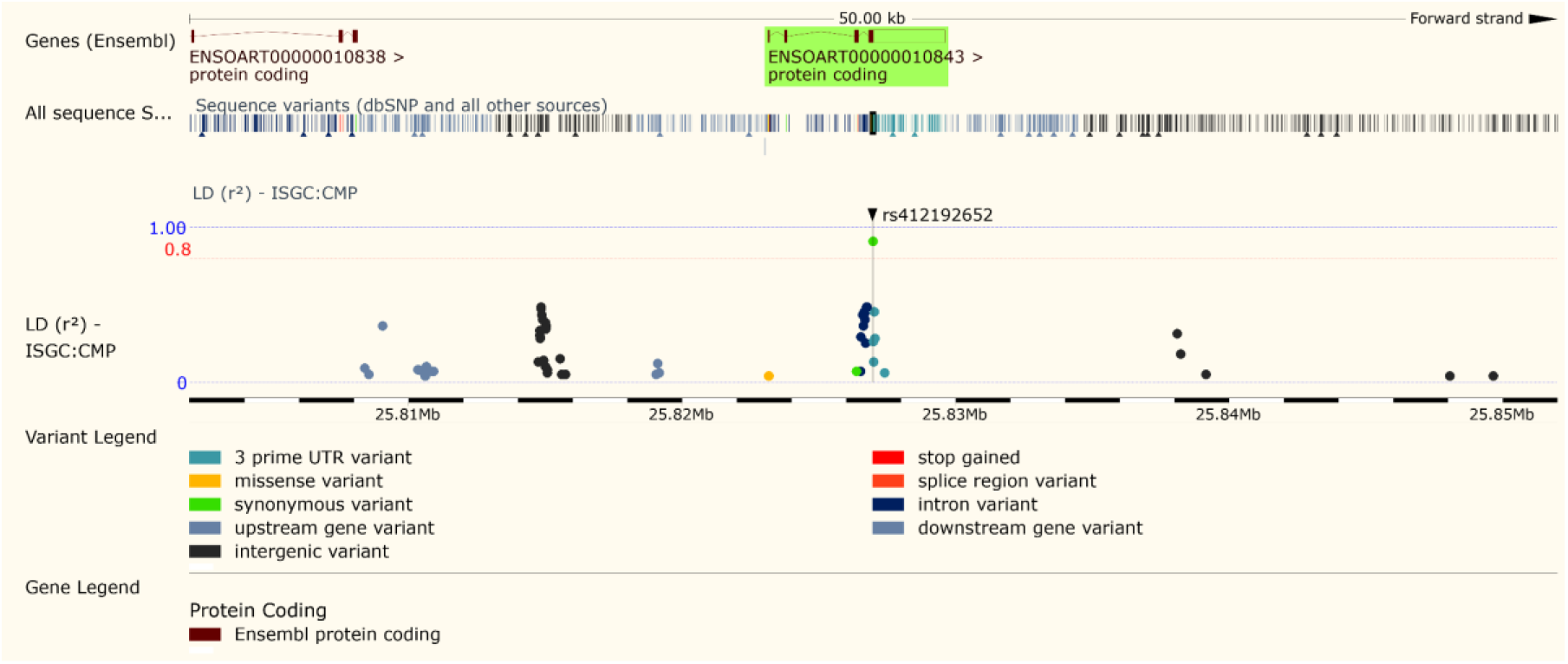
The LD block (A) and LD Manhattan plot (B) of SAA3 transcript (ENSOART00000010843). The LD block produced as part of International Sheep Genome Consortium is publicly available on Ensembl using the COMPOSITE population (n = 98).

**Figure S5.**
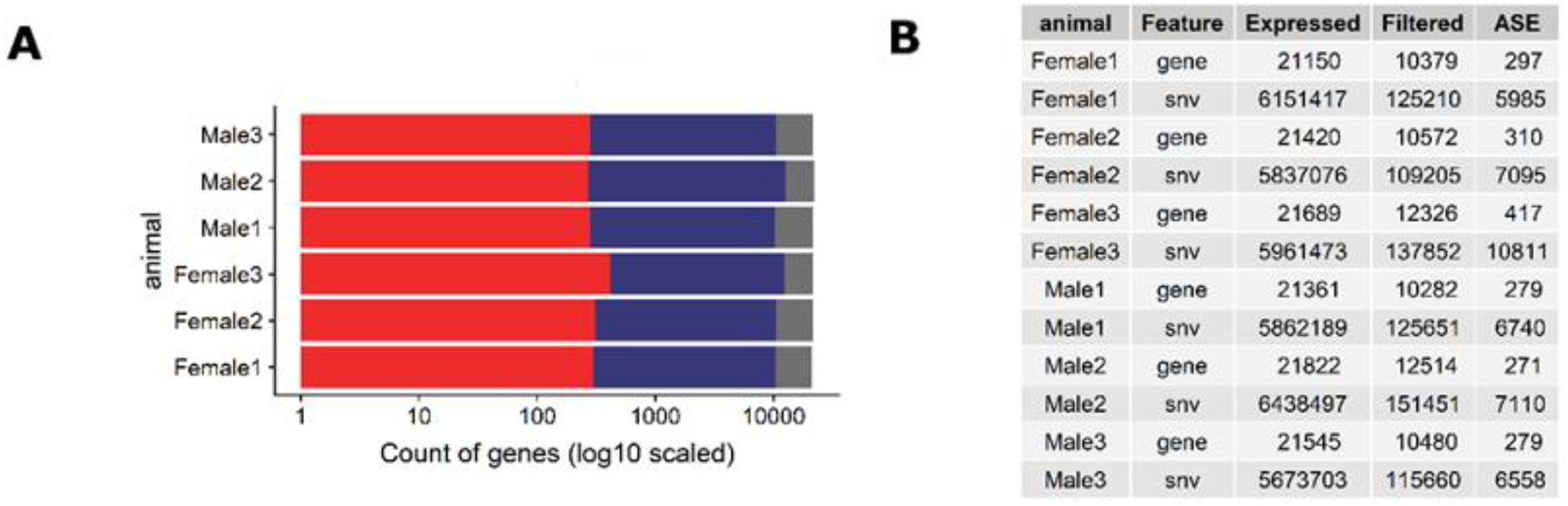
The proportional distribution of ASE genes and SNVs being expressed across tissues averaged for each sheep. A) Count of genes in 6 sheep expressed in grey, informative/filtered in blue and ASE positive in red. B) Break down of genes and SNVs that are expressed, pass minimum filtration criteria and are ASE positive.

**Figure S6.**
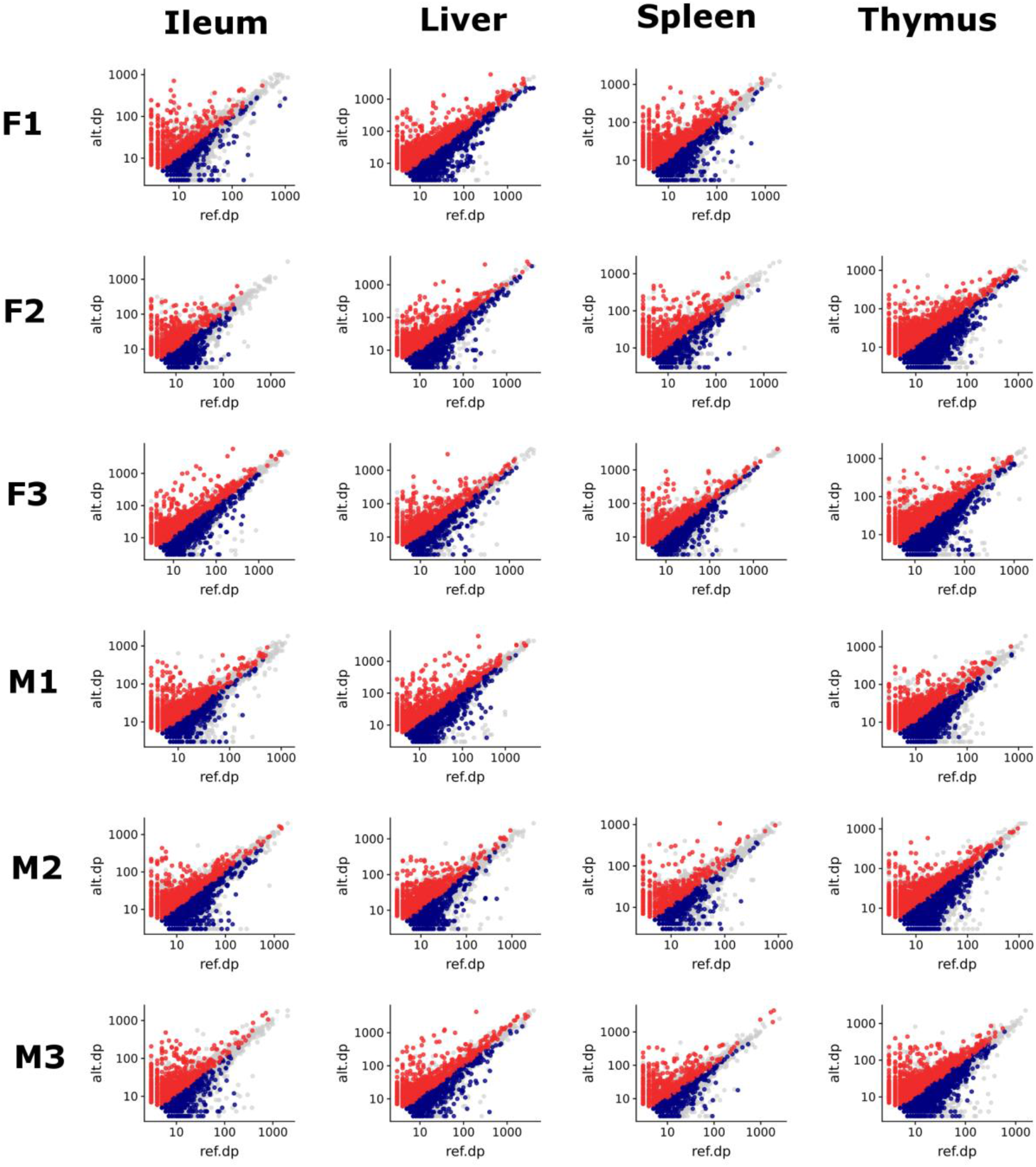
Scatter plot of allelic depth (read count) in all 4 tissues of the 6 TxBF sheep. The SNVs passing minimum filtration criteria are coloured in grey and significant ASE (FDR10%) are marked by red and blue. If reference read count > alternate read count the point is shown in blue and vice versa in red. Expression level varies across tissues but does not affect the ASE profile.

**Figure S7.**
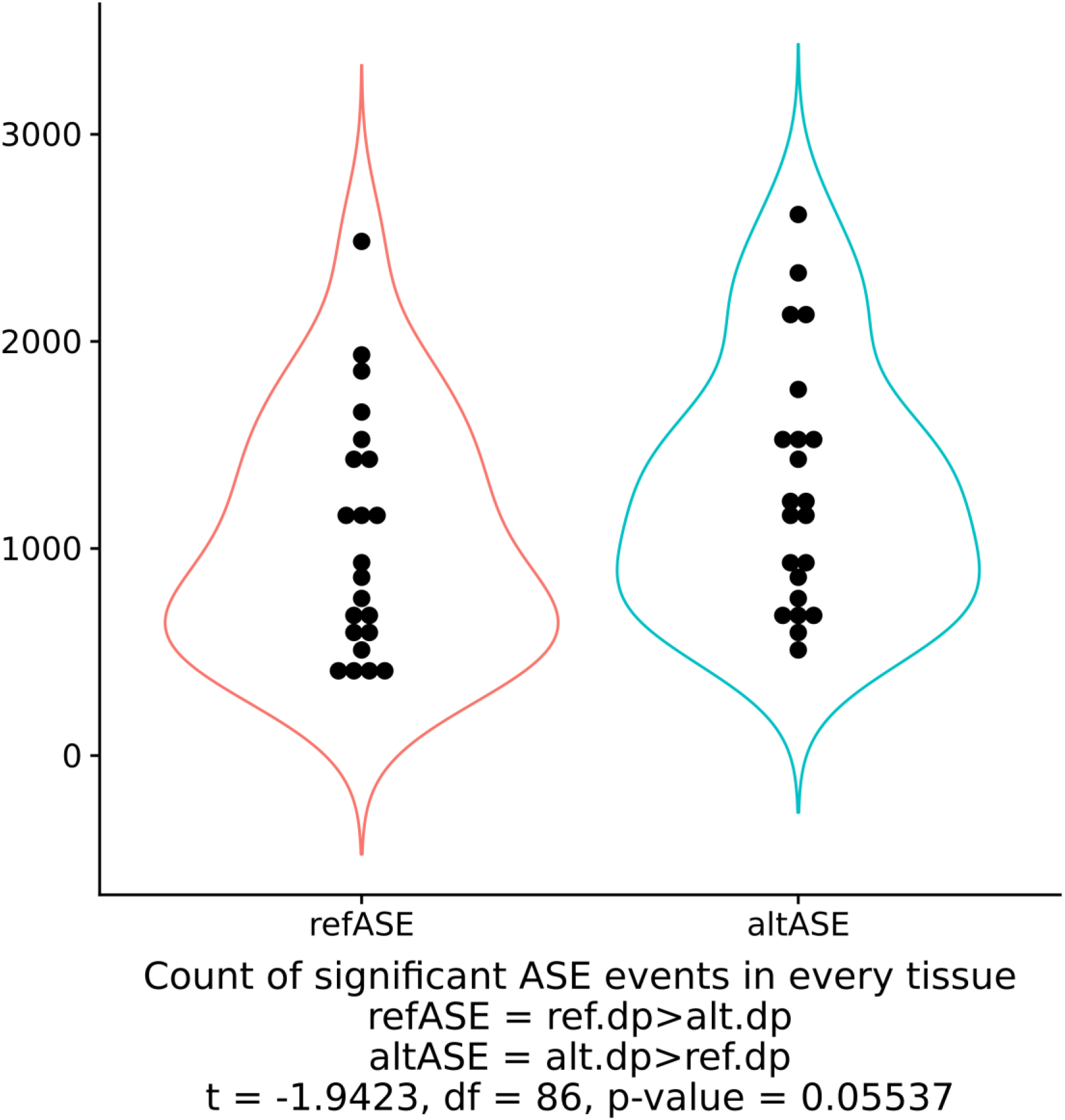
Distribution of ASE positive SNVs over the direction of imbalance in the 34 tissues of 6 sheep. Ref.dp > Alt.dp as ASE towards Ref allele and vice versa. There was no significant difference between the count of ASE events given the direction of the allelic imbalance (p-value > 0.05).

**Figure S8.**
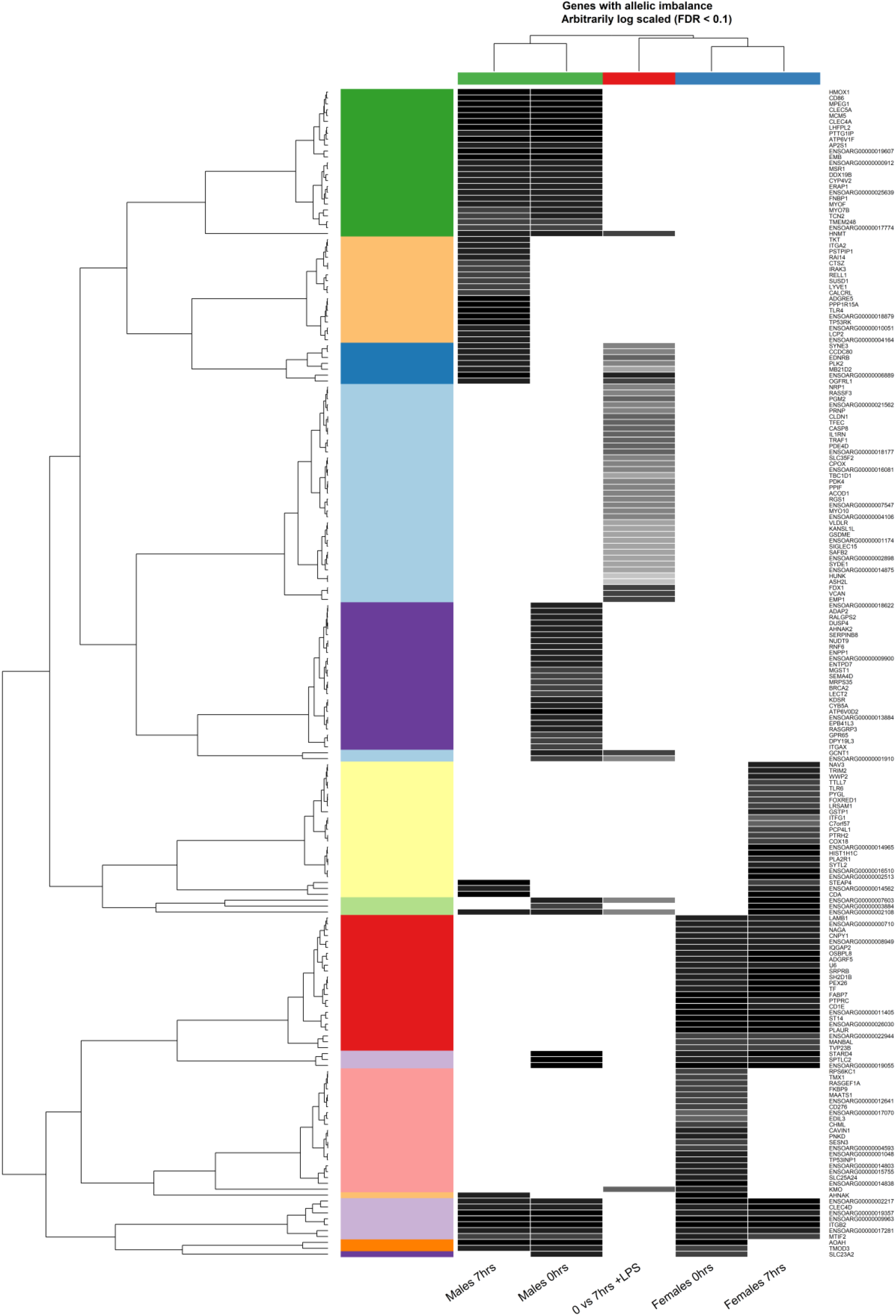
The Kmeans cluster analysis of the ASE genes in between static and ICD profiles of BMDMs at 0 and 7hrs samples. Heat map dataframe has been log and centre scaled prior to cluster analysis. The heatmap shows the ASE genes in 5 groups: 0vs 7 hrs ICD set, Males 0hrs, Males 7hrs. Females 0hrs and Females 7hrs.

**Figure S9.**
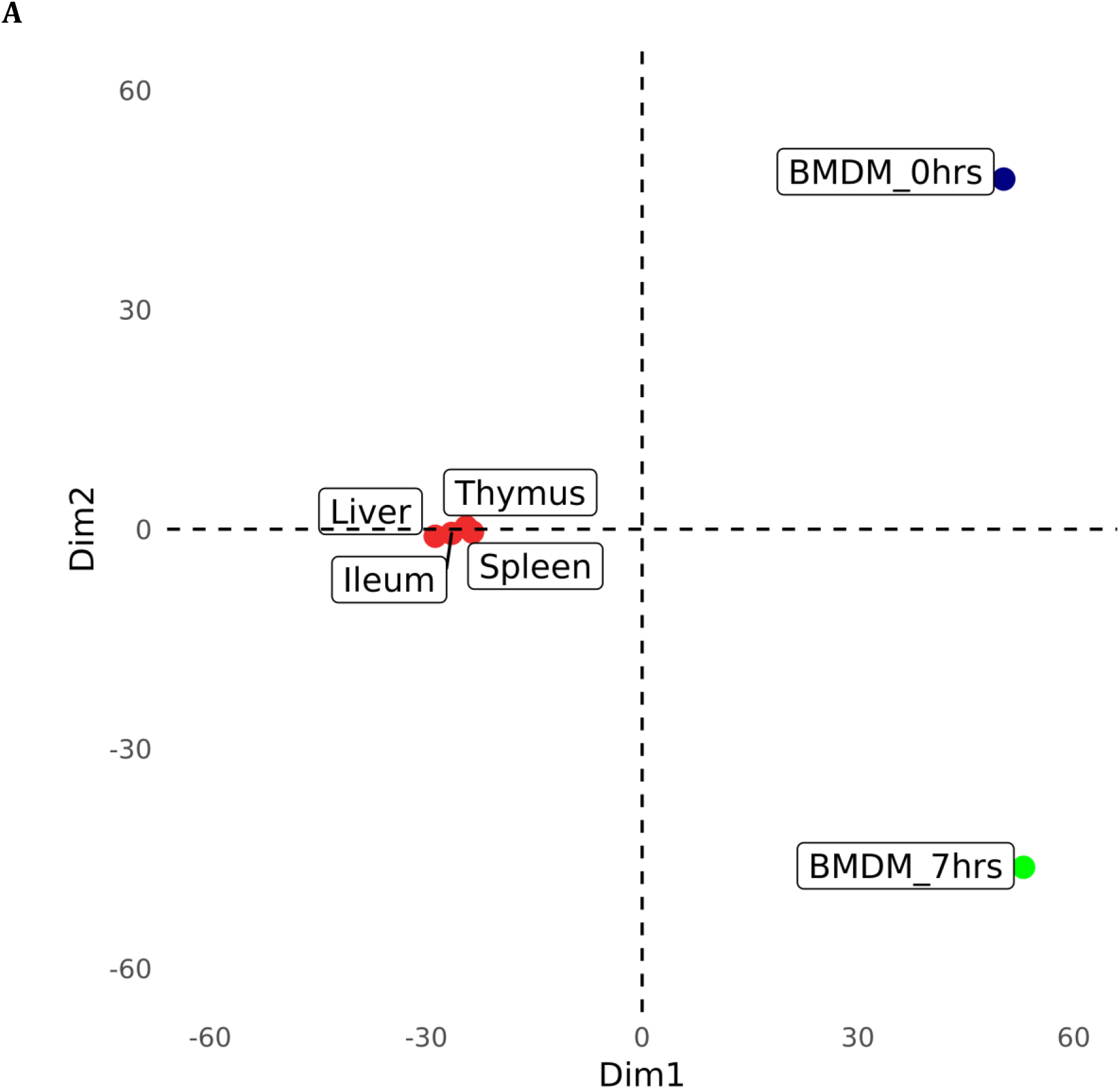
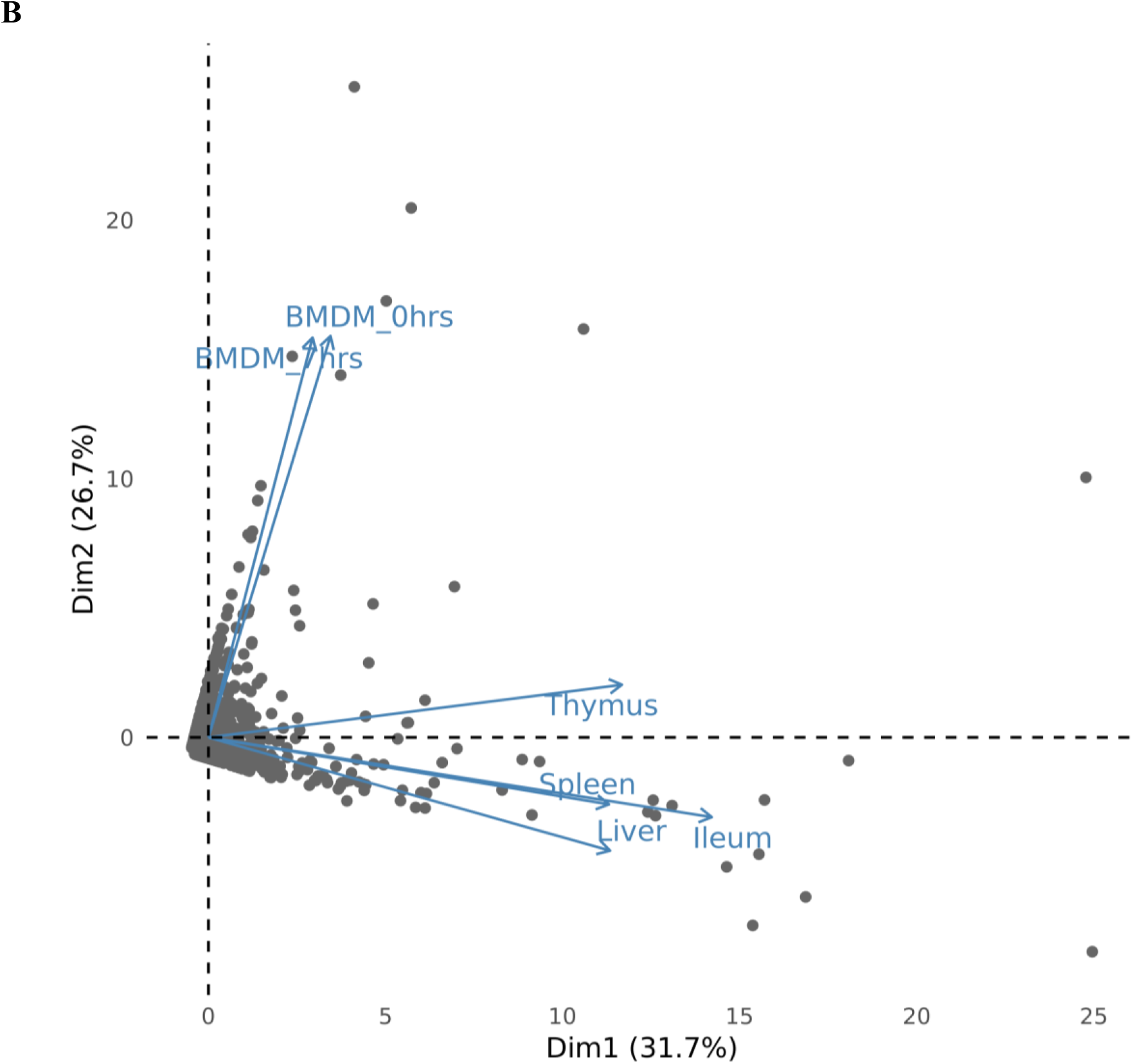
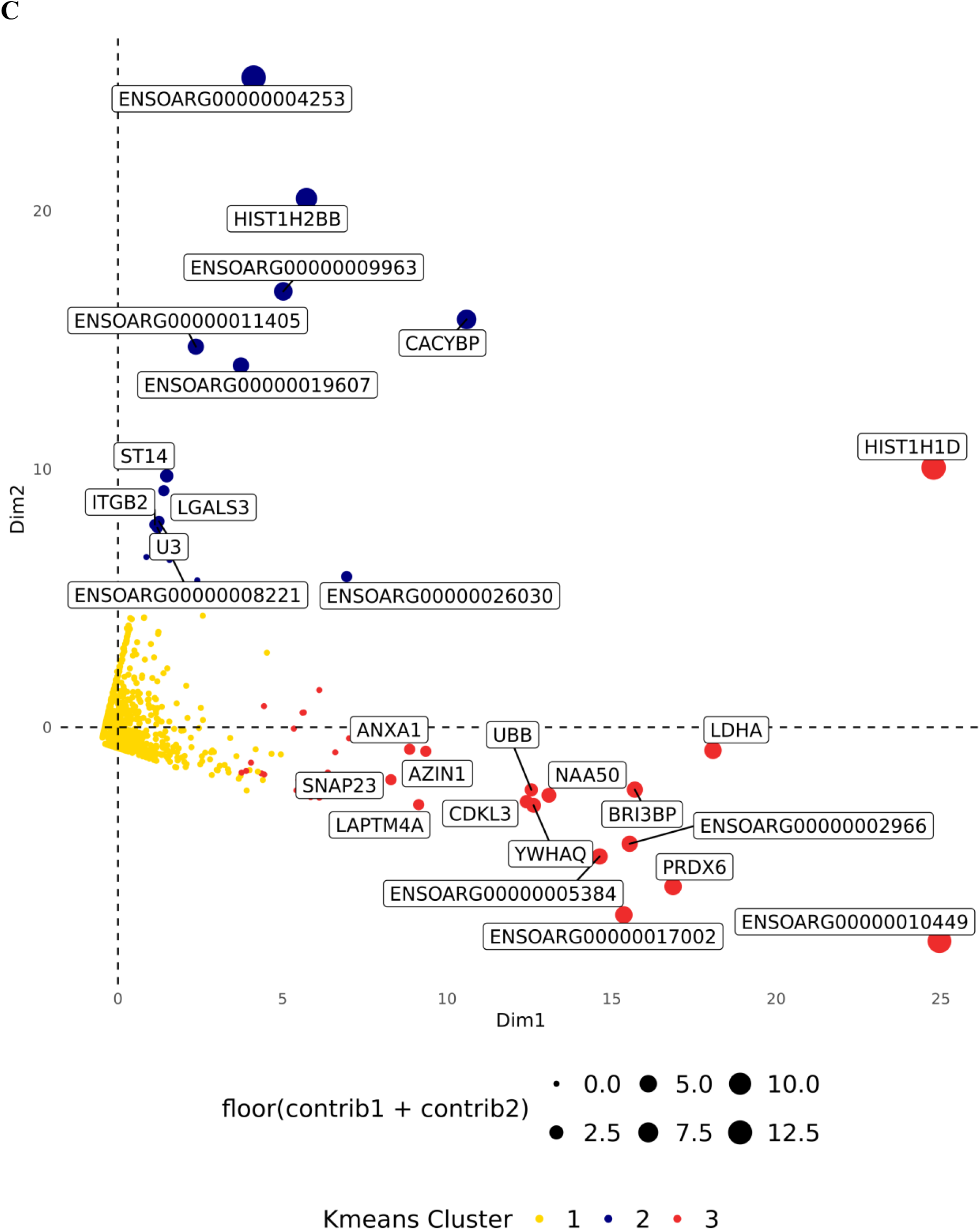
Principle component analysis and Kmeans clustering of ASE gene profiles of 4 tissues and 1 cell type (2 treatment) in 6 sheep. The gene list comprised of all ASE genes in each tissue or cell type was used to perform PCA which established the presence of a hierarchical stratification resulting in 3 groups in the dataset. This grouping was used to cluster the dataset using K means algorithm with 3 centres. **A)** Samples PCA plot of the first 2 PCs (Matrix 6 sample rows x 3000 gene columns) **B)** Genes PCA plot of the first 2 PC (Matrix 3000 gene rows x 6 sample columns) **C)** Kmeans Clusters overlay on the PC1 vs PC2 scatter plot from section B (Genes PCA plot). Gene name labels were included only for the genes with overall contribution to the first 2 components > 1.

**Figure S10.**
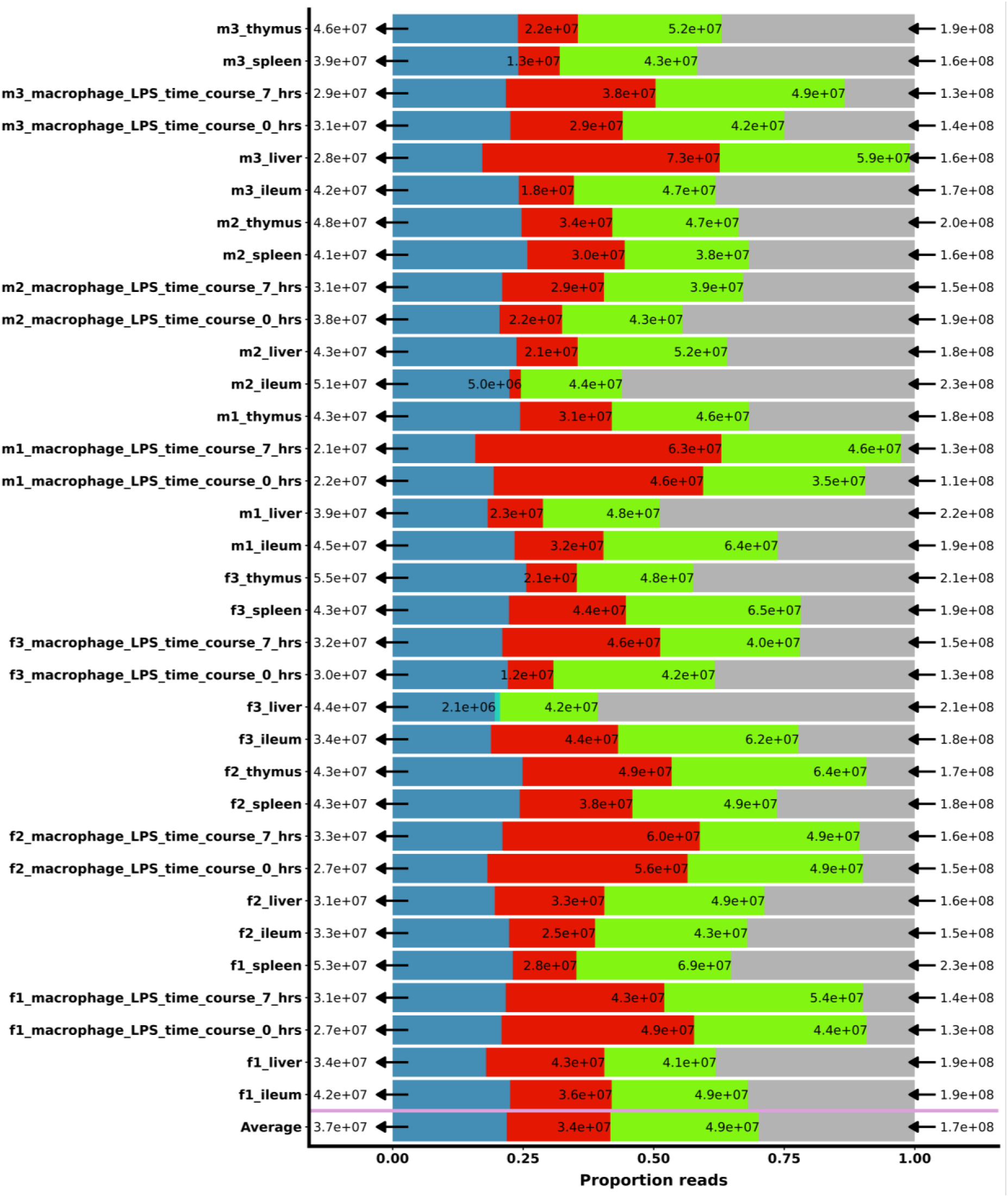
The read loss proportion during the WASP reference mapping bias removal steps. The graph shows the input (read counts on the right) and output (read counts on the 2^nd^ left) RNA-Seq reads from the study before and after WASP processing respectively. Each bar represents 4 overlaid read proportions labelled by colour and raw read count. The total portion of reads coloured in grey shows 100% of the reads processed by WASP (input 170 million reads on average). The green portion shows the non-intersecting reads (and their duplicates) mapped within the range of Ensemble gene coordinate boundaries (exonic, intronic and UTRs reads 49 million reads on average). The red portion of the reads represents reads intersecting bi-allelic heterozygote sites and selected for synthetic read production step (34 million reads on average). The blue portion of the graph represents the final output of WASP’s pipeline (reference biased and duplicated reads removed from red and green portions respectively). The output portion (37 million reads on average) was used in GATK ASEReadCounter for allelic read counts. The last row on the bottom shows the average of these proportions across both tissue and cell type samples. Reference biased reads count for average 20% of input RNA-Seq data.

**Figure S11.**
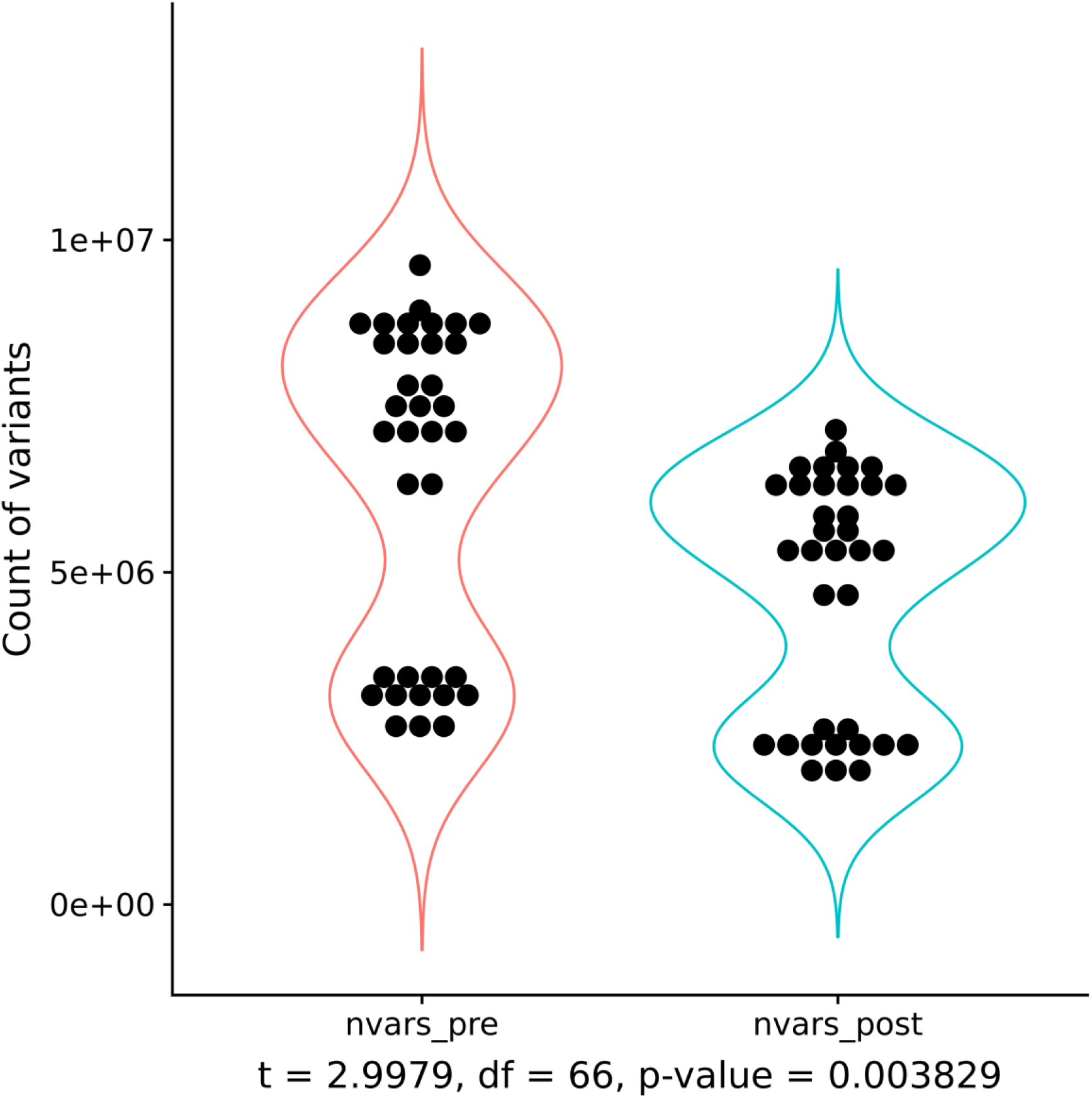
Number of SNVs before and after WASP reference bias correction. There is a significant reduction in number of variants captured (25% on average 1.5e+07) between two steps. Tissues samples dots mainly present within the upper quantile and BMDMs in the lower. Average number of variants beforehand = 6,325,570 and afterwards = 4,729,639.

**Figure S12.**
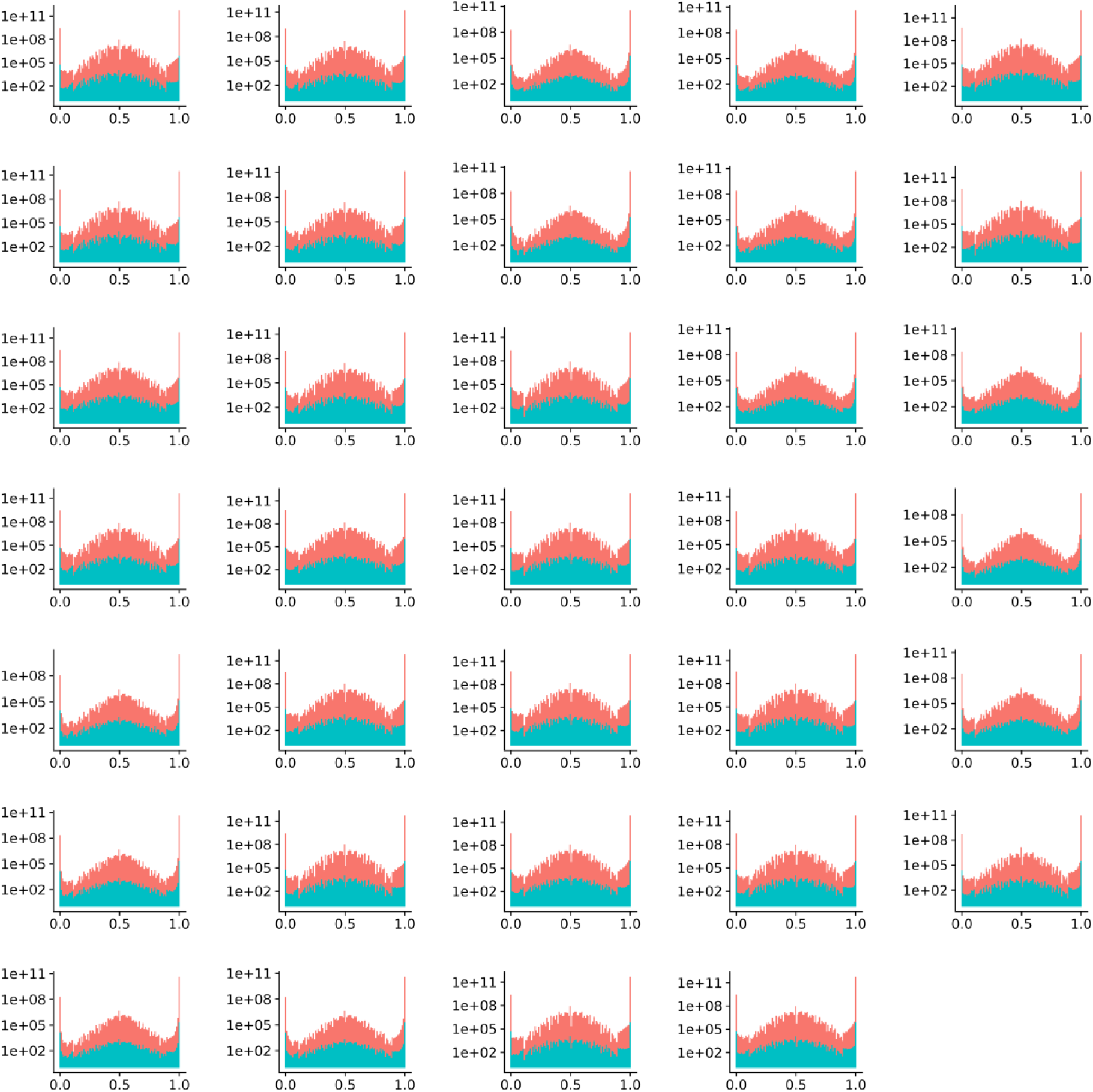
The comparison of reference allele count ratio before and after WASP mapping bias correction. The distribution of refRatio (refCount/refCount+altcounts) in all the tissue and cell samples before (in red) and after WASP (in green). The overall counts, left skew (*viz*. range 0.75-1) and refRatio=1 inflation is significantly reduced after WASP bias correction in all samples.

**Table S1.**
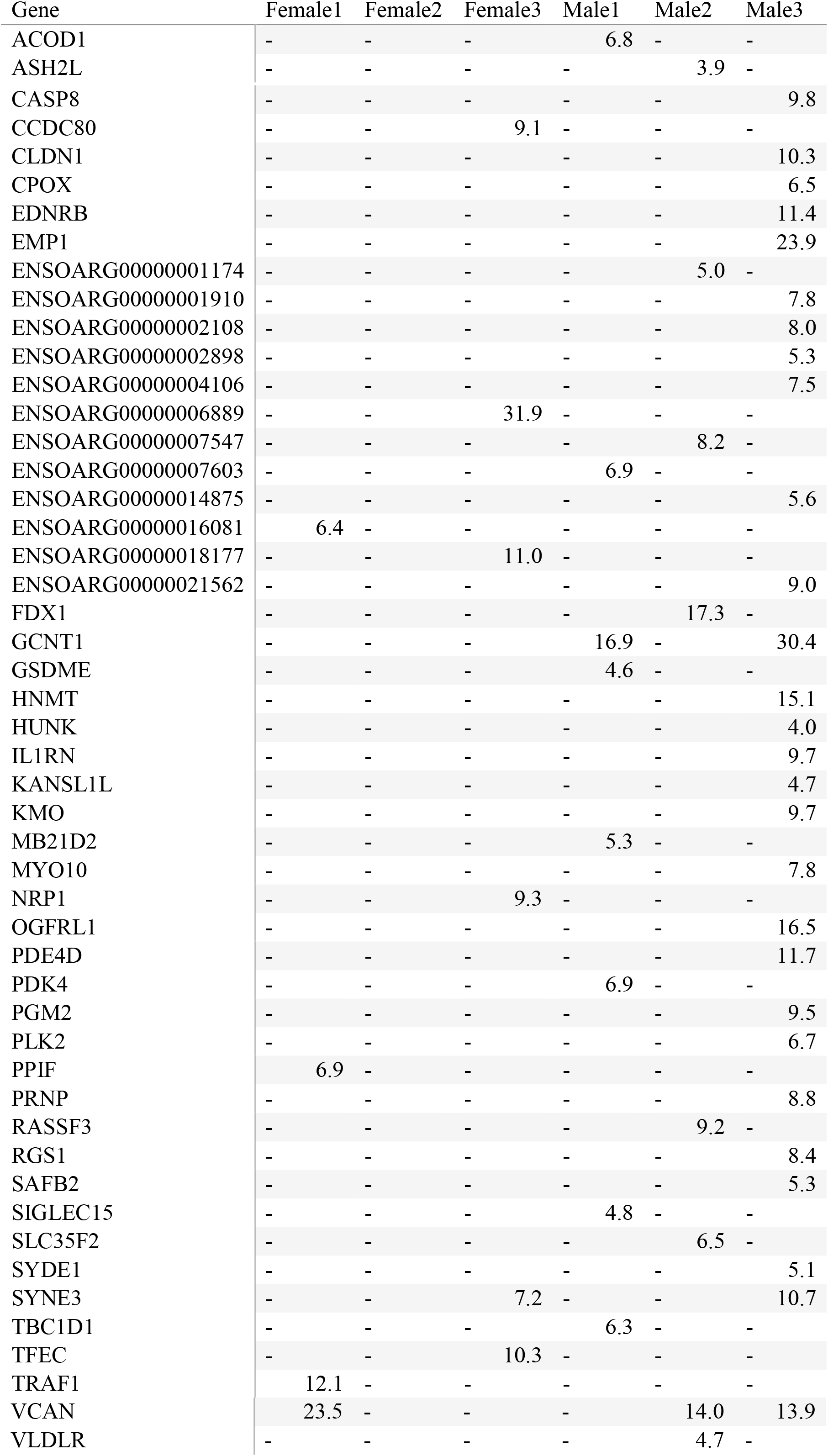
Mean allelic imbalance (log2(ASE_+LPS_/ASE _-LPS_)) in all 6 sheep. Only 1 gene (VCAN including 7 SNVs) exhibited inducible ASE which was shared between 3 sheep (in bold), while two (GCNT1 and SYNE3) showed inducible ASE in two sheep. Diverse and highly individual ASE response to LPS was observed in each sheep.

